# Multi-omics analysis identifies loci associated with pyrethroid resistance across sister species in the *Anopheles gambiae* species complex

**DOI:** 10.64898/2026.01.30.702793

**Authors:** Juan Carlos Lol, Antonia L Böhmert, Juliane Hartke, Antoine Sanou, Jakob Kerbl-Knapp, Marion Morris, Julia B Mäurer, Patrick Hogan, Moussa Wamdaogo Guelbeogo, Eric R Lucas, Hilary Ranson, Victoria A Ingham

## Abstract

The *Anopheles gambiae* species complex includes *An. gambiae, An. coluzzii*, and *An. arabiensis*, three of the four major malaria vector species across Africa. Insecticide-based vector control remains the most critical tool in the fight against malaria, with a heavy reliance on pyrethroid insecticides. However, widespread pyrethroid resistance jeopardises the effectiveness of these methods. Currently, the number of diagnostic markers available to track pyrethroid resistance in endemic settings is limited, focusing mainly on changes at the insecticide target site. Genetic analysis of seven insecticide-resistant populations from West Africa, a region with intense insecticide resistance, reveals shared loci associated with pyrethroid resistance among the sister species of the *An. gambiae* complex including *kdr, GSTE1-8* and the *CYP6P* region. Notably, a mutation in the voltage-gated sodium channel, I1527T, previously identified in *An. coluzzii* and *An. gambiae*, was detected in wild-caught *An. arabiensis* from Burkina Faso. Additionally, a mutation, L207I, in *GSTE7* common to *An. coluzzii* and *An. arabiensis*, significantly increases survivorship to deltamethrin. With the additional integration of RNAseq and data from the *Anopheles gambiae* 1000 genomes project, we were able to identify putative eQTLs associated with the expression of major insecticide resistance-related transcripts, such as *CYP6P3, CYP9K1* and *CYP6AA1*. This study reveals several potential diagnostic markers of resistance that can be implemented in endemic settings and identifies putative introgression between *An. arabiensis* and *An. coluzzii* in Burkina Faso.

## Introduction

Malaria remains one of the world’s most deadly diseases, with over 280 million cases and around 610 000 deaths in 2024, predominantly in African children (WHO 2025).Malaria is caused by the *Plasmodium* parasite and is transmitted to humans through the bite of infected *Anopheles* mosquitoes. In sub-Saharan Africa, four major malaria vectors account for the majority of the transmission: *An. funestus* and three sibling species: *An. gambiae, An. coluzzii* and *An. arabiensis* (Sinka, et al. 2010; Coetzee, et al. 2013). These mosquitoes differ in their biting behaviour and ecological niches but all display a preference for night-time biting (Kabbale, et al. 2013; Sangbakembi-Ngounou, et al. 2022; Nzioki, et al. 2023), making insecticide treated bed nets (ITNs) the most effective way to control malaria transmission. Since 2004, nearly three billion ITNs, all treated with pyrethroid insecticides, have been distributed (WHO 2025). Pyrethroids target the nervous system by interacting with the voltage-gated sodium channel, causing continuous activation, leading to paralysis and death. The widespread use of ITNs and agrochemicals have imposed intense selective pressure on *Anopheles* mosquitoes, resulting in ubiquitous pyrethroid resistance across Africa (Sadia, et al. 2024). Indeed, the intensity of resistance is so high in West Africa, that these mosquitoes can contact pyrethroids multiple times with no impact on their longevity (Hughes, et al. 2020).

Although pyrethroid resistance is a complex trait (Ingham 2023), involving multiple mechanisms within the same populations, mechanistic convergence is seen across diverse species (Ingham and Nagi 2024). For example, mutations in the voltage-gated sodium channel, the target site of pyrethroids, known as knockdown resistance (*kdr*) are well-characterized, with four prominent variants associated with resistance: L995F/S (Martinez-Torres, et al. 1998; Ranson, et al. 2000) and V402L-I1527T (Clarkson, et al. 2021). L995F, L995S and V402L-I1527T are mutually exclusive (Clarkson, et al. 2021), with the former conferring higher levels of pyrethroid resistance than the latter in transgenic mosquitoes; however, L995F has an associated fitness cost (Grigoraki, et al. 2021; Williams, Cowlishaw, et al. 2022). L995 mutations are found across the *Anopheles gambiae* species complex, whilst V402L-I1527T has only been detected in *An. coluzzii* (Clarkson, et al. 2021; Kientega 2023) and *An. gambiae* (Collins, et al. 2019) and appears to be replacing L995F in West Africa (Williams, Cowlishaw, et al. 2022; Kientega 2023). Analogously, metabolic resistance resulting from overexpression of enzymes, such as cytochrome P450s, UGTs, ABC transporters and glutathione-S-transferases (GSTs) is seen across all major *Anopheles* species (Ingham, et al. 2018; Ingham and Nagi 2024). Cytochrome P450s have been shown to directly bind to and metabolise pyrethroids (Yunta, et al. 2016; Yunta, et al. 2019; Nolden, et al. 2022), an ABC transporter has been shown to reduce uptake of pyrethroids (Kefi, et al. 2023), inhibition of UGTs restores susceptibility to pyrethroids (Logan, et al. 2024), whilst GSTs have been indirectly linked with pyrethroid resistance (Riveron, et al. 2014; Opondo, et al. 2016; Ingham, Bennett, et al. 2020; Tao, et al. 2022; Ibrahim, et al. 2023). Several less well-characterised mechanisms have been linked to pyrethroid resistance, including cuticular thickening (Balabanidou, et al. 2016), sequestration (Isaacs, et al. 2018; Ingham, Anthousi, et al. 2020; Ingham, Brown, et al. 2021) and behavioural changes (Killeen and Chitnis 2014).

Tracking the spread of insecticide resistance and understanding the underlying molecular mechanisms are crucial for informing the future use of vector control tools and understanding the spread of resistance. Although the role of specific metabolic enzymes in insecticide resistance has been clearly delineated, genetic markers to track this resistance remain largely elusive. The *Anopheles gambiae* 1000 genomes project (Ag1000G) was conceived to provide a high-resolution overview of genetic diversity of major malaria vectors across multiple countries (Anopheles gambiae Genomes, et al. 2017; Anopheles gambiae Genomes 2020) and has generated phased haplotypes from over 25,000 wild-caught mosquitoes covering 31 African countries (Consortium 2021). Ag1000G has been used to identify copy number variations (CNVs) in the *An. gambiae* complex associated with resistance and molecular diagnostics have been developed to track these mutations (Lucas, Miles, et al. 2019; Lucas, Rockett, et al. 2019; Lucas, et al. 2023). CNVs in the *CYP6AA/CYP6P* locus were shown to be increasing in frequency in the Democratic Republic of the Congo, Kenya and Uganda (Njoroge, et al. 2022) and are associated with resistance to deltamethrin in Côte D’Ivoire (Kouame, et al. 2023) and to permethrin and the insecticide synergist PBO (Mawejje, et al. 2013) in Uganda. In addition to this, duplicated SNPs associated with *CYP9K1* (Mawejje, et al. 2013), a carboxylesterase (Weetman, et al. 2018) and *GSTE2* (Opondo, et al. 2016; Lucas, Rockett, et al. 2019; Diallo, et al. 2022) have been used to track the spread of pyrethroid resistance-related alleles across multiple countries. However, for many resistance genes with the clearest association with pyrethroid resistance, including *CYP6P3* and *CYP6M2* no diagnostic markers are available despite distinct haplotypes associated with resistance (Lucas, et al. 2023). Several other regions, likely related to immune response in the mosquito such as Thioester-containing protein 1 (*TEP1*) have also shown to be under selection (White, et al. 2011), with putative indication of involvement in insecticide resistance (Ibrahim, et al. 2023; Lucas, et al. 2023).

The selection pressure induced by insecticide use has led to adaptive introgression of multiple resistance-related loci. For example, 995F-*kdr* introgressed from *An. gambiae* to *An. coluzzii* (Clarkson, et al. 2014) and resistant alleles of the target site of dieldrin (*Rdl*) introgressed from *An. gambiae* and *An. arabiensis* to *An. coluzzii* (Ingham, Tennessen, et al. 2021). However, the degree to which metabolic resistance associated mutations have introgressed, or arisen *de novo* in different species is largely unknown.

In this study we perform whole genome sequencing on five colony populations encompassing the three major vector species in the *An. gambiae* species complex and field-caught *An. coluzzii* from the same locality. We additionally integrated WGS from three colonies from a previous publication (Ingham, Tennessen, et al. 2021). Our findings reveal an intricate evolutionary history involving shared loci across sister species of the *An. gambiae* complex. We demonstrate that a sweep in the antiparasitic *TEP1* is not associated with insecticide resistance, and for the first time, we report the presence of the I1527T mutation in both colony and wild-caught *An. arabiensis*. Additionally, we identify a novel mutation in *GSTE7*, L207I, linked to deltamethrin survival. Alongside whole-genome sequencing, RNAseq data for each colony allowed for broad cis-eQTL analysis. When integrated with allele and sweep data from Ag1000G, this analysis revealed several putative SNPs that may drive the increased expression of insecticide resistance-related loci.

## Results

### Variant calling

Individual mosquitoes from five colonised resistant populations and one wild-caught population of the *An. gambiae* species complex underwent whole genome sequencing (n=110; Supplementary Table 1). A further two resistant colonies (30 individuals total) and a susceptible colony (20 individuals) sequenced in previous work were included (Ingham, Tennessen, et al. 2021), totalling 160 mosquito samples. One colony, Tiassalé_Ag.sl. (*An. gambiae* s.l.), was included as a colony outgroup both due to uncertainty in species and its origin in Côte D’Ivoire, whilst all others originate in Burkina Faso. The species represented are *An. gambiae, An. coluzzii*, and *An. arabiensis* and are described in Table 1 and prior publications (Williams, et al. 2019; Williams, Ingham, et al. 2022). Of the 160 samples, three were removed for unexpectedly out-grouping within the PCA analysis, leaving 157 samples grouping as expected (Figure 1A). In total 12,389,338 high-quality variants were detected across the 157 samples, of which 9,615,152 were biallelic; the chromosomal breakdown is described in Table 1. Nucleotide diversity was unsurprisingly highest in the Tengrela (F) field population (*An. coluzzii*), although all colony populations retain diversity across autosomes (Supplementary Figure 1A). The lab populations showed high levels of Tajima’s D, reflecting the sudden population contraction associated with lab colonisation (Supplementary Figure 1B).

**Table 1:**
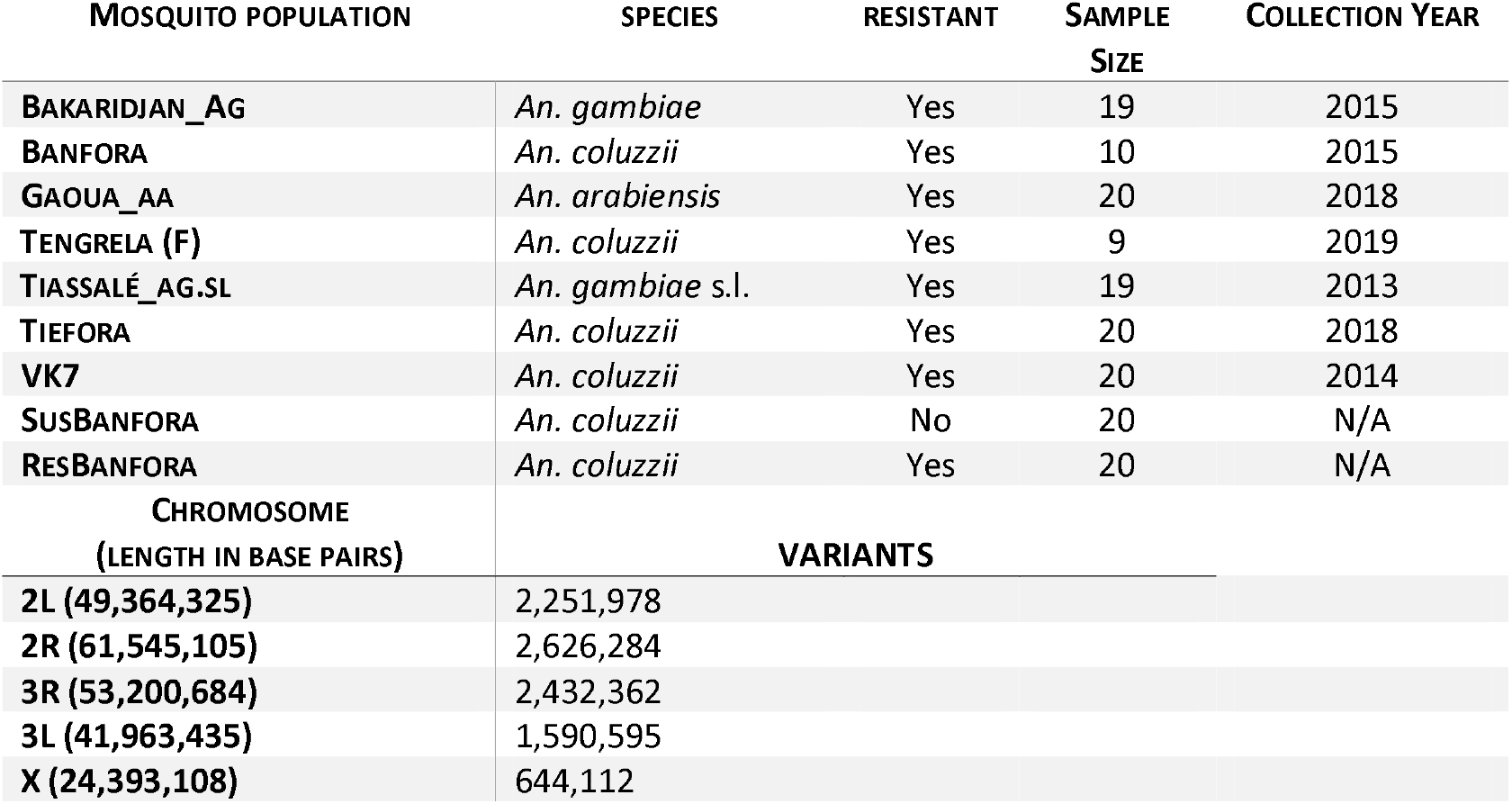
Sample breakdown and called high quality, biallelic variants. Sample names contain reference to species: Ag = *An. gambiae*, Aa = *An. arabiensis* and Ag.sl = *An. gambiae* s.l., all others are *An. coluzzii*. (F) denotes the field population. The final column indicates the date of collection, with the exception of Tengrela (F), all populations were colonised and maintained under insectary conditions prior to WGS.

SNPEff predictions indicated that of the biallelic SNPs, 2,451 (0.025 %) represent high impact changes and 270,264 (2.81 %) with moderate impact (Supplementary Table 2). Within the high impact SNPs several relate to the chemosensory protein cluster, recently implicated in pyrethroid resistance (Ingham, Anthousi, et al. 2020). Here, SNPEff predictions show *SAP3* (AGAP008054) is differentially spliced in *An. arabiensis* compared to the other members of the species complex, whilst *CSP1* (AGAP008059) demonstrates the same differential splicing within Tiassalé_Ag.sl, Tiefora (*An. coluzzii*) and Gaoua_Aa (*An. arabiensis*) colonies. Within the metabolic enzymes linked with insecticide resistance, only *CYP6Z4* is impacted, with a stop codon gain resulting in loss of the terminal amino acid in Tengrela (F) and Bakaridjan_Ag (*An. gambiae*) populations.

### Population Structure

For all population genetics analysis SusBanfora and ResBanfora were excluded, being derived from the original Banfora colony. The *An. gambiae* species complex phylogeny has been resolved showing ((*An. gambiae, An. coluzzii*), *An. arabiensis*) (Thawornwattana, et al. 2018); however, both SplitsTrees and unrooted maximum likelihood trees on chromosome 3L show ((*An. gambiae, An. arabiensis*), *An. coluzzii*) relationship within these populations from Burkina Faso (Figure 1B, 1C). Whilst SplitsTrees supports this when Tiassalé_Ag.sl is included, this population outgroups in an unrooted maximum likelihood phylogeny. SplitsTrees displays clear evidence of connectivity between the different species, indicative of an interwoven evolutionary history, which is absent in the *An. coluzzii* Tiefora population. Within the *An. coluzzii* grouping in both SplitsTrees and maximum likelihood phylogeny, Banfora unexpectedly groups with VK7, which were collected over 100 km away. Both the Tiefora and Tengrela (F) populations, although temporally disparate, were collected respectively at 10 km and 25 km from Banfora, indicating complexity within the *An. coluzzii* species. To further assess evolutionary history, admixture analysis was carried out. As expected, the majority of colonies show single ancestry due to population bottlenecking; however, a single *An. arabiensis* individual appears to show a fraction of common ancestry with the *An. coluzzii* population from Tiefora (Figure 1D). Interestingly, the field-caught *An. coluzzii* Tengrela (F) population displays mixed ancestry across all colonised populations, with a very small fraction from *An. arabiensis* and a slightly larger fraction from *An. gambiae* reflecting expected phylogenetic relationships.

**Figure 1:**
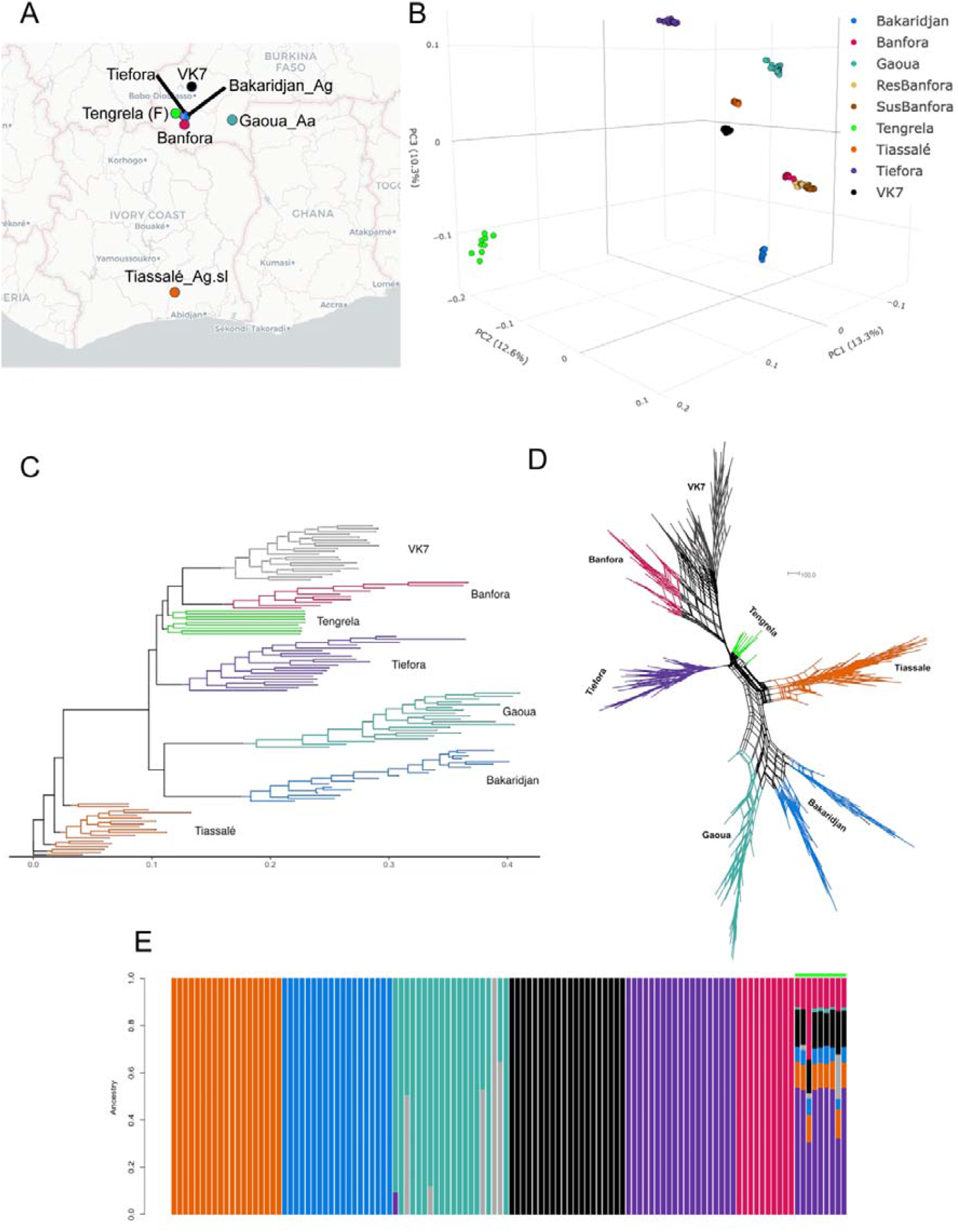
Phylogenetics and population structure. **A**. Collection sites of populations with the non-*An. coluzzii* populations labelled as in Table 1: Ag = *An. gambiae*, Aa = *An. arabiensis* and Ag.sl = *An. gambiae* s.l., all others are *An. coluzzii*. (F) indicates the field population. Map produced using CartoDB through MapView (https://carto.com/) **B**. 3D PCA plot showing the whole genome PCA for all nine populations (157 samples) analysed in this study, PC1 (x), PC2 (y) and PC3 (z) shown and labelled with variance attributed to each component. **C**. Unrooted maximum likelihood phylogeny based on chromosome 3L based on 10,000 bootstraps, branch length indicates number of changes per site, with each population coloured: VK7 (grey), Banfora (red), Tengrela (F) (green), Tiefora (purple), Gaoua_Aa (teal), Bakaridjan_Ag (blue) and Tiassalé_Ag.sl (orange). **D**. Consensus SplitsTrees network with evolutionary distance indicated based on 10,000 bootstraps. Populations are coloured as in C. **E**. STRUCTURE analysis of inferred ancestry for each population, based on k = 7. Populations coloured as previously; grey indicates variation in *An. arabiensis* and green indicates variation in Tengrela (F).

### Inversion status

Mosquitoes of the *An. gambiae* complex have large chromosomal inversions across chromosome 2 (Love, et al. 2020). The inversion statuses of the populations were determined using previously published tagging SNPs (Figure 2A) (Love, et al. 2020). The largest inversion, 2La, was fixed in all populations except the *An. gambiae* s.l. population, Tiassalé_Ag.sl, which had 9 individuals heterozygous. Of the 2R inversions: 2Rb was at high frequency in Bakaridjan_Ag and Gaoua_Aa and intermediate frequency in all populations with the exception of Tiassalé_Ag.sl; 2Rc was at intermediate frequency in Tengrela (F) and Tiefora, in addition to one VK7 individual and 2Ru was found at high frequency in Tiefora and intermediate frequencies in Tengrela (F), Banfora and VK7.

**Figure 2:**
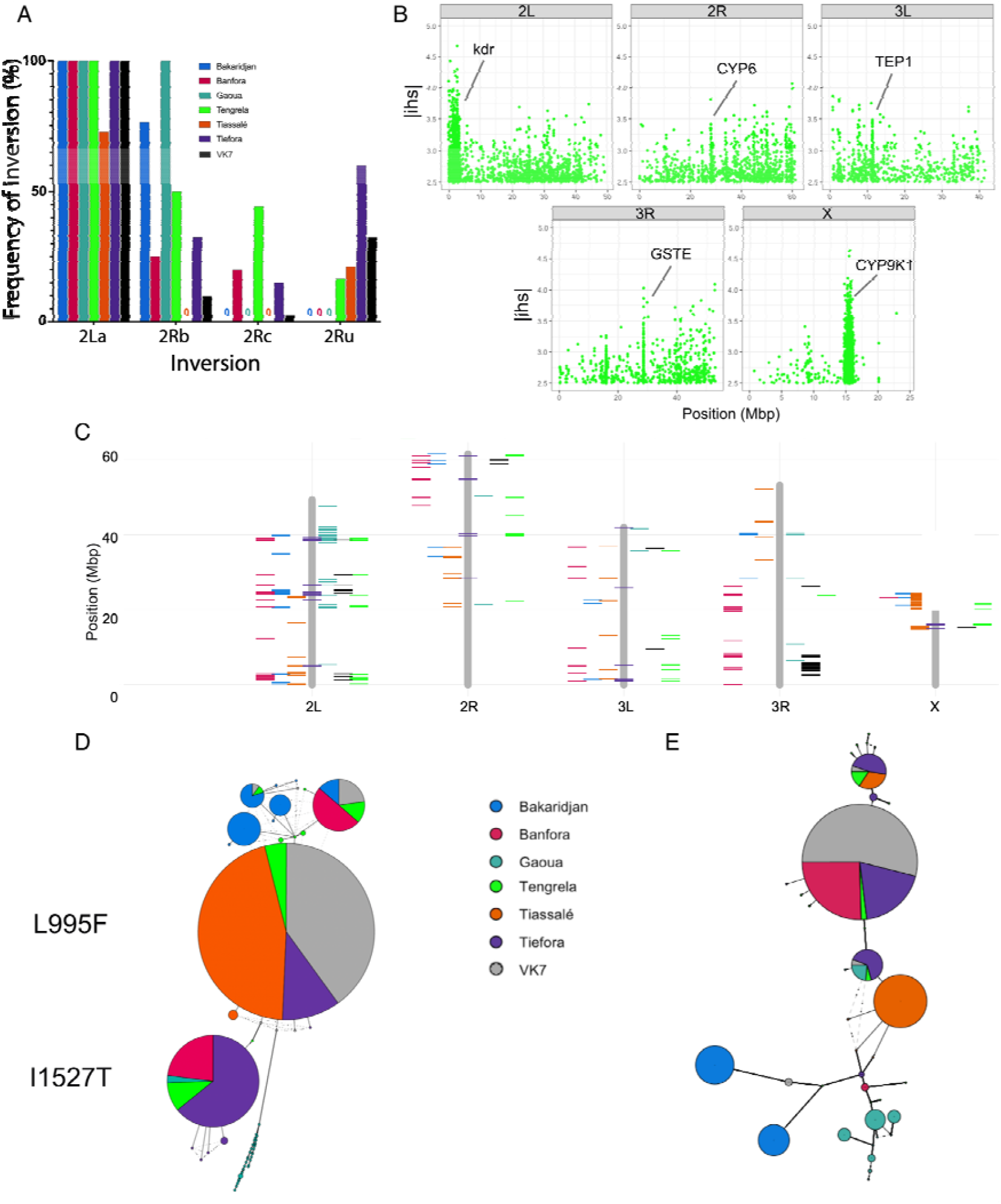
Patterns of genome wide selection. **A**. Chromosomal inversion frequencies in each population, coloured by population. **B**. Genome-wide integrated haplotype statistic for the field caught Tengrela (F) population. Peaks around loci putatively linked with immunity and insecticide resistance are labelled. **C**. Top 5 % of the overlaps of FST and ABBA-BABBA statistics for each population, with positions (y-axis) illustrated across each individual chromosome (x-axis). Populations coloured as in A. **D**. Haplotype map of the 10 kb region surrounding the *kdr* locus. Clear splits between the L995F and I1527T haplotype are labelled. **E**. Haplotype map of the 10 kb region surrounding the *GSTE* loci. Both **D**. and **E**. show in-grouping of *An. arabiensis* within *An. coluzzii* haplotypes, indicative of putative introgression.

### Selective sweeps

iHS and iHH12 (H12) statistics were generated for each population to define areas of extended haplotype homozygosity and thus inferred selection (Voight, et al. 2006; Garud, et al. 2015). Signatures of extended haplotype homozygosity were taken as the overlap of the top 1 % of the absolute calculated scores for each statistic (Supplementary Figure 2, Supplementary Figure 3). Tengrela (F), a wild caught *An. coluzzii* population demonstrated clear selective sweeps around resistance-related loci, including *kdr, CYP9K1* the *GSTE* cluster and the *CYP6P* cluster (Figure 2B, Supplementary Figure 4). Interestingly, the peak H12 SNP in the *GSTE* locus centres on I114T, a known non-synonymous mutation in GSTE2 conferring DDT resistance (Mitchell, et al. 2014), whilst in *CYP9K1* it centres on a synonymous SNP (Supplementary Figure 5). Although this region is often attributed to CYP9K1, there are a total of 49 genes within the sweep and the iHS peak corresponds to an intragenic region upstream of AGAP000821 (unknown function). Overall, 14 clear blocks of selection are seen in Tengrela (F) and whilst five can be attributed to resistance-related loci, the others largely overlap with previous sweeps identified in Ag1000G including the large block at the end of 2R, also seen in Ag1000G (Anopheles gambiae Genomes, et al. 2017). A peak around the *TEP1* locus is evident, the major immune response mediator to *Plasmodium* (White, et al. 2011) and previously linked to insecticide resistance (Ibrahim, et al. 2023; Lucas, et al. 2023).

Similar analysis was performed on the lab colonies (Supplementary Figure 2, Supplementary Figure 3), with the caveat that they have undergone a large bottleneck and will have strong selection not only for insecticide resistance due to being maintained under consistent selection pressure but for membrane feeding and survival under insectary conditions. Amongst the loci under strong selection and known to be involved in resistance within the colony populations was *Keap1*, which showed strong signatures of selection in *An. gambiae* and *An. arabiensis* populations and some evidence of selection in both VK7 and Tiefora *An. coluzzii* populations. Both the shape of the sweep and its predominance in *An. gambiae* matches that seen in Ag1000G (Nagi 2025). Interestingly, the *An. gambiae* and *An. arabiensis* populations share a selective sweep around the *GSTD* locus, which is also present in Tiefora. There is no evidence of a sweep around the *kdr* locus in any of colonies. The strength and intensities of the iHS and H12 statistics varies across the colonies, likely reflecting time since the bottlenecking of the colony, and variations in colony size over time (Supplementary Figure 2, Supplementary Figure 3).

### Shared Haplotypes

Firstly, to rule out contamination across species, X-chromosome ancestry informative markers were taken for Ag1000G and all species were as expected (Supplementary Figure 6). To determine potential areas of introgression, shared haplotypes were determined between the different species of the *An. gambiae* complex (Figure 2C; Supplementary Figure 7). Shared haplotypes are defined here as loci that fall within the top 5% of the windowed ABBA–BABA statistics (Durand, et al. 2011) and bottom 5% of diversity corrected F_ST_ values in the paired introgressed population. The relationship between introgression signal strength and genetic differentiation was assessed by quantifying the genome-wide association between the absolute estimated fraction of introgressed ancestry |fdM| and F_ST_ for each focal population using Pearson’s and Spearman’s correlation coefficients. Across focal populations, the genome-wide association between |fdM| and F_ST_ was generally weak and in all cases negative, consistent with the expectation that windows with stronger introgression signals can exhibit reduced differentiation. The magnitude and direction of this relationship varied among populations, reflecting differences in evolutionary history and the extent of introgression. Permutation tests identified significant enrichment of donor-specific low-F_ST_ supported fdM outliers in some populations, whereas others showed no deviation from the null expectation (Supplementary Table 3), indicating heterogeneous introgression across populations rather than a uniform genome-wide process.

Within the overlap, 1128 genes were within 10 kb of these regions (Supplementary Table 4) and enriched in GO terms for structural component of the cuticle (2.07e-14) and marginally non-significant for monooxygenase activity (6.37e-2). Whilst caution is needed interpreting shared haplotypes from colonised populations, it is interesting that these regions include *CYP9K1*, the *CYP6P* region, *CYP4G16* the *CSP* cluster, a number of cuticular clusters and *Rdl* (Table 2; Supplementary Figure 8A-E), which have all been linked to resistance (Ffrench-Constant, et al. 1993; Edi, et al. 2014; Balabanidou, et al. 2016; Yunta, et al. 2016; Yunta, et al. 2019; Ingham, Anthousi, et al. 2020; Khan 2020). *CYP9K1* shows a similar haplotype across *An. gambiae* and *An. coluzzii* whilst the CYP6P locus showed shared haplotypes between Tiassalé_Ag.sl, Tiefora, VK7 and one Tengrela (F) individual, in addition to one Banfora individual displaying an *An. arabiensis*-like block (Supplementary Figure 8C). *CYP4G16* shows species-specific differences, with *An. arabiensis* being more variable. Both the *CSP* cluster and Rdl show high levels of diversity, with the CSPs having distinct overlap between Tiefora and *An. arabiensis*.

**Table 2:** Binary presence or absence table for the region being within the top 5% of |fdM| values. Region of interest and sample names, where ‘1’ indicates a |fdM| within the top 5% and ‘0’ indicated |fdM| outside this range.

| Region of interest | Bakaridjan.Ag | Gaoua.Aa | Tengrela (F) | Tiassale_AG.sl | Tiefora | VK7 | Banfora |
| --- | --- | --- | --- | --- | --- | --- | --- |
| CYP6P cluster | 0 | 0 | 0 | 1 | 1 | 1 | 0 |
| CYP4G16 | 0 | 0 | 0 | 1 | 0 | 0 | 0 |
| CYP9K1 | 0 | 0 | 1 | 1 | 1 | 1 | 1 |
| GSTE cluster | 0 | 0 | 1 | 0 | 1 | 1 | 1 |
| CSP cluster | 0 | 0 | 0 | 0 | 0 | 1 | 1 |
| kdr | 1 | 0 | 1 | 1 | 0 | 0 | 0 |
| Rdl | 1 | 1 | 1 | 0 | 1 | 1 | 1 |
| TEP1 | 1 | 1 | 1 | 1 | 1 | 1 | 0 |

Next, regions strongly associated with selective sweeps observed in field populations (Nagi 2025) were explored for introgression in an *a priori* manner using only the |fdM| values. Here, *kdr* (Figure 2D), *GSTE1-8* (Figure 2E), *COEAE1F/2F, Keap1* and *Ace1* were explored and of these only *kdr* and *GSTE1-8* demonstrated |fdM| within the top 5% (Table 2). Interestingly, on closer inspection of these regions through haplotype clustering, potential haplotype sharing between *An. arabiensis* and *An. coluzzii* in the *kdr* (Figure 2D) and *GSTE* loci (Figure 2E) was evident. Two individual *An. arabiensis* showed the I1527T *kdr* mutation on the same genetic background as previously reported in Banfora and Tiefora and also seen here in wild caught Tengrela (F). Exploration of the non-synonymous SNPs found within the *GSTE* cluster showed that four SNPs are located within this block. In *An. arabiensis*, three individuals were heterozygous and one homozygous for the four non-synonymous SNPs, two in *GSTE1* and two in *GSTE7*. All four of these were also present in Tiassalé_Ag.sl, Tiefora, Tengrela (F) and VK7. The mutations in I114T and L119V within *GSTE2*, which have been widely associated with resistance (Riveron, et al. 2014; Lucas, Rockett, et al. 2019; Diallo, et al. 2022) were present in these populations. Specifically, I114T was found in Tiassalé_Ag.sl and across all *An. coluzzii*, as previously reported (Williams, Ingham, et al. 2022), and L119V was found in *An. gambiae* and one Tengrela (F) sample.

### *Kdr* locus

Shared haplotypes were evident between two *An. arabiensis* individuals and several *An. coluzzii*, corresponding to the V402L-I1527T mutation, currently spreading through *An. coluzzii* in Burkina Faso [16]. V402L-I1527T has never been reported in *An. arabiensis*, and so to confirm the WGS data, a TaqMan assay was used, which verified the presence of this allele at low frequency (<5 %) within the Gaoua_Aa *An. arabiensis* colony across multiple generations (G8-G54). To determine whether this mutation is also found in wild caught *An. arabiensis* from the same region, and thus rule out contamination, the TaqMan assay was performed on *An. arabiensis* previously collected from two regions of Burkina Faso in 2021 and 2022 (Supplementary Table 5). Of the 95 samples, six failed, four were positive for the I1527T mutation and the remainder were wildtype. Two of the four positive samples were collected in Tiefora in July-August 2021 and two in Ouagadougou (Zogona) in November 2022. Of these, the two samples from Tiefora were sequenced and confirmed as heterozygotes. Further exploration of the *kdr* locus in *An. arabiensis* revealed a previously unreported non-synonymous mutation at A1553T, which is present at a frequency of 45 % in this colony. Mutation A1553T is located in the domain III of the voltage-gated sodium channel where other *kdr* mutations have been found associated with insecticide resistance in mosquitoes (Nagi 2025). A mutation at position 1553 was also reported in Helicoverpa armigera and Heliothis virescens potentially conferring pyrethroid resistance. Non-synonymous substitutions in the *kdr* locus in each population are described in Table 3.

**Table 3:**
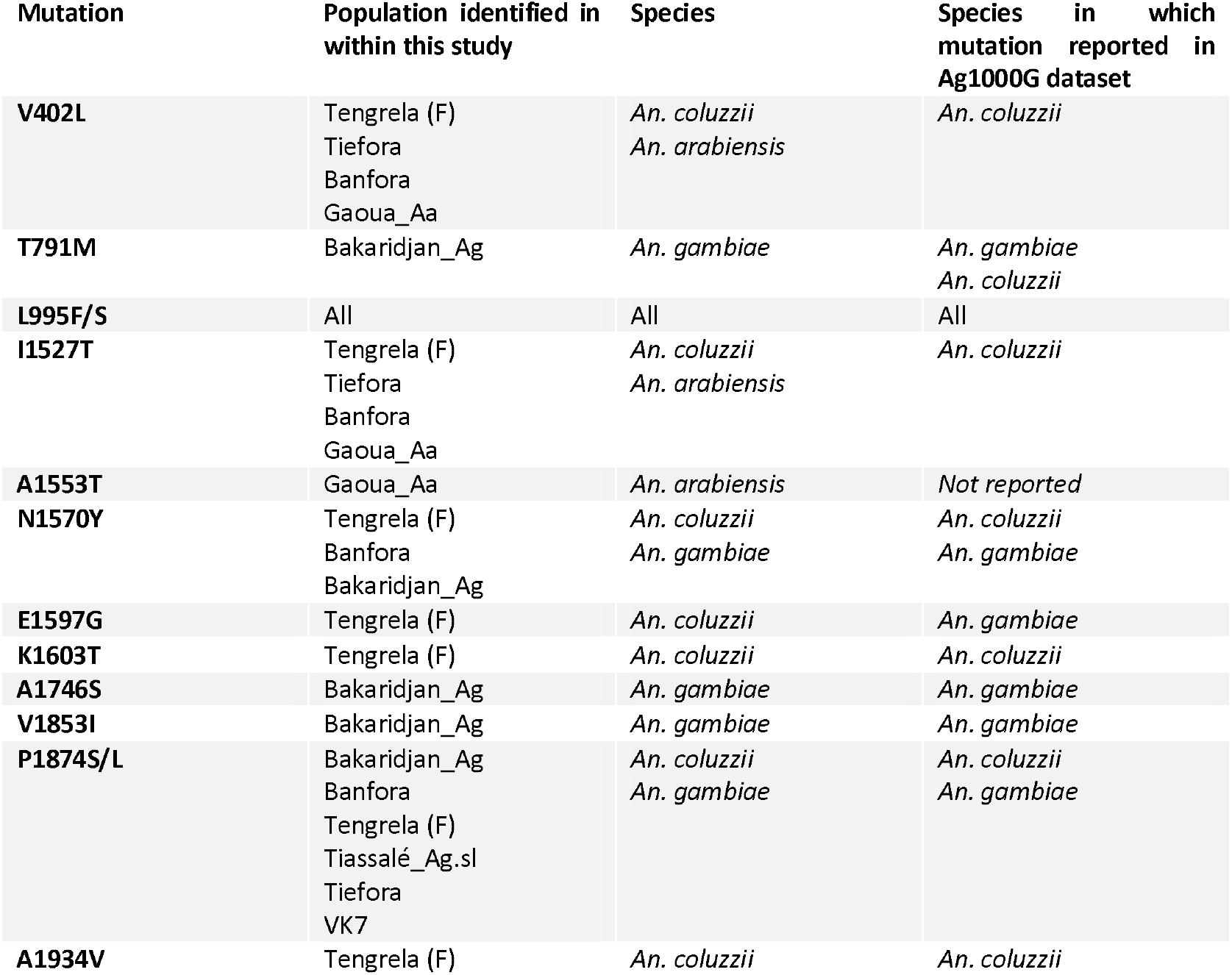
Architecture of the *kdr* locus. Mutations identified in this dataset, the populations in which these are present, corresponding species and presence in the Ag1000G dataset [58]. Ag = *An. gambiae*, Aa = *An. arabiensis* and Ag.sl = *An. gambiae* s.l., all others are *An. coluzzii*. (F) shows the field population.

### GSTE loci

GSTs in the Epsilon family have previously been reported to be involved in resistance to DDT and associated with resistance to pyrethroids (Ortelli, et al. 2003; Riveron, et al. 2014) and so this locus was explored in more detail (Figure 3A). Examination of the loci highlighted four non-synonymous SNPs associated with potential introgression between *An. coluzzii* and *An. arabiensis* in *GSTE1* (AGAP009195) and *GSTE7* (AGAP009196) (L37F/E83D and P3R/L207I respectively) with two heterozygote samples and one sample fixed for the mutations. Next, population of *An. arabiensis*, 88-field caught samples previously screened for the *kdr* allele were then assayed by PCR amplification and sanger sequencing. Unexpectedly, 43 failed and the other samples showed the wild-type allele for the L207I mutation (Supplementary Table 5). To determine whether L207I mutation contributes to insecticide resistance in the *An. gambiae* complex, an allelic discrimination PCR was designed that distinguishes L and I at position 207 in *GSTE7* in mosquitoes. Tiefora and Tiassalé_Ag.sl were used to determine whether there was an association of mutation L207I with survival after exposure to 5X diagnostic dose (DD) of deltamethrin, alpha-cypermethrin and permethrin through WHO tube assays (Supplementary Figure 9A, B). A clear association was seen with deltamethrin survivorship and the L207I mutation in *An. coluzzii* (Fisher’s Exact Test, *P* < 0.0001, Figure 3B), but not with permethrin or alpha-cypermethrin (Supplementary Figure 9 C, D). The comparison of the different genotypes showed a significant association of the heterozygous genotype (Fisher’s Exact Test, *P* < 0.0001) with deltamethrin survivorship. Similarly, in the case of *An. gambiae* s.l. (Figure 3C, Supplementary Figure 9 E, F), no association of the L207I mutation was seen with permethrin or alpha-cypermethrin but borderline statistical non-significance was found with deltamethrin (Fisher’s Exact Test, *P* = 0.0571). The comparison of the different genotypes showed a significant association with survival when heterozygous, as with *An. coluzzii* (Fisher’s Exact Test, *P* = 0.0252), indicating a survival advantage for heterozygous mosquitoes in both species.

**Figure 3:**
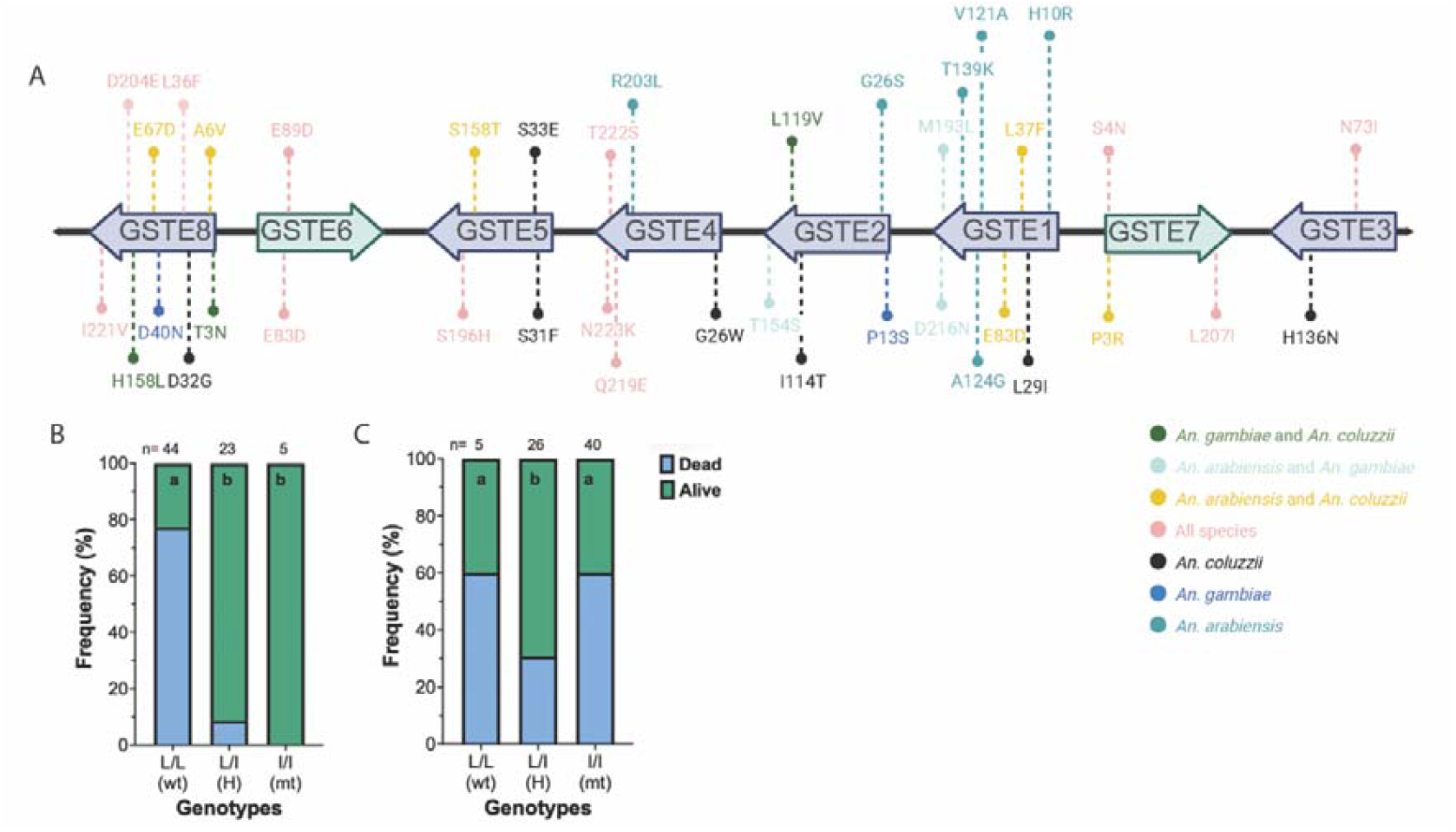
Mutations found in the GSTE locus. **A**. Schematic diagram of the GSTE locus with non-synonymous SNPs found in the populations highlighted. Mutations are represented by a dotted line, and the associated non-synonymous mutation is shown above. Colours correspond to the key below. Reverse strand genes are shown in green. **B. C**. Association study of L207I with resistance to the 5X DD using WHO tube tests of three pyrethroids in the **B**. Tiefora and **C**. Tiassalé_Ag.sl populations. n shows number of mosquitoes screened, wt = wild-type, H = heterozygous, mt = mutant, significance calculated by Fisher’s Exact Test and different letters in the bars indicate significant differences in the proportion of genotypes in deltamethrin-resistant mosquitoes.

### *TEP1* locus

*TEP1* is associated with a selective sweep in the field population Tengrela (F) (Figure 2B) and has previously been linked with selection in Ag1000G (Consortium 2021). The *Plasmodium* resistant *TEP1* allele is nearly fixed in *An. coluzzii* from Burkina Faso (Tennessen, et al. 2021), and the gene has been shown to be upregulated in multiple insecticide resistant populations (Ibrahim, et al. 2023; Ingham and Nagi 2024) and linked to resistance (Lucas, et al. 2023). To investigate any role in pyrethroid resistance in driving this sweep, dsTEP1 injection was used to significantly reduce transcript expression compared to dsGFP controls (78 % reduction 72 hours post injection (hpi) and 77 % 96 hpi compared to control, Supplementary Figure 10A). Protein expression was similarly reduced at 72 hpi, with 75 % reduction in processed *TEP1* and complete absence of full length *TEP1* (Supplementary Figure 10B). Reduction in transcript and protein levels led to no change to mosquito longevity (Supplementary Figure 10C). Mosquitoes were then exposed to 0.05 % deltamethrin 72 hpi, and no significant difference in mortality was seen between dsGFP and dsTEP1, suggesting this sweep is likely caused by selection pressure due to *Plasmodium* infection and is unrelated to insecticide resistance (Supplementary Figure 10D).

### Copy Number Variation

Copy number variants (CNVs) were identified in each newly sequenced population using previously published methodologies (Lucas, Miles, et al. 2019) (Figure 2F, Supplementary Table 6). In the *An. gambiae* population (Bakaridjan_Ag), CNVs include the previously reported duplication of *CYP9K1* (Cyp9k1_Dup11) and a potential duplication event near the cuticular protein of low complexity (*CPLC*) cluster on chromosome 3R. In the *An. arabiensis* population (Gaoua_Aa), duplications were observed around *GSTE1, GSTE2* (Gstue_Dup12), *TEP1*, and *CYP6Z4*, as well as near *CPLC* genes on chromosomes 3L and 3R. Amongst *An. coluzzii* populations, the field-caught Tengrela (F) samples exhibited duplications involving *CYP6AA1, CYP6AA2* (Cyp6aap_Dup1b, Cyp6aap_Dup7 and Cyp6aap_Dup10), *TEP1*, and the CPLC cluster on chromosome 3R. The Tiefora population showed duplications in *CYP6AA1, CYP6AA2* (Cyp6aap_Dup7 and Cyp6aap_Dup10), GSTE2 (Gstue_Dup1), *TEP1, APL1C, COEAE6O*, and the *CPLC* cluster on chromosome 3R. Similarly, the VK7 population displayed duplications in *CYP6AA1, CYP6AA2* (Cyp6aap_Dup1b, Cyp6aap_Dup7 and Cyp6aap_Dup14), *TEP1, HEX1A* (a hexamerin), *GSTE2* (Gstue_Dup1), *COEAE6O*, and the *CPLC* cluster on chromosome 3R. Finally, in the *An. gambiae* s.l. population from Tiassalé_Ag.sl, CNVs were observed near *TEP1* and the *CPLC* cluster on chromosome 3R. Across the identified CNVs, several genes were consistently observed in multiple populations, including heat shock proteins (AGAP004581-4583), vitellogenin, the *CYP325* family, and various ionotropic and gustatory receptors.

### eQTL identification

Pooled RNAseq data was available for seven of the sequenced colony populations (Ingham, Tennessen, et al. 2021; Williams, Ingham, et al. 2022), whilst a new RNAseq dataset was produced for Tiassalé_Ag.sl. The count files from these experiments were then used to simulate individual read counts. A pseudo-expression quantitative trait locus (eQTL) analysis was performed using the simulated counts based on pooled RNAseq and identified a total of 1,069,779 SNPs significantly associated with gene expression. To filter these results to relevant transcripts, genes previously associated with insecticide resistance, including detoxification-related proteins, neuronal targets, and chemosensory proteins (listed in Supplementary Table 7), were used. Out of the 343 genes contained in this list, 304 were associated with one or more eQTLs (Supplementary Table 8), totalling 41,920 SNPs. The large number of putative associated SNPs is attributed to extensive haplotype blocks due to low resolution of this dataset. Next, to determine whether these SNPs are relevant in field populations, their presence in field-caught *An. gambiae* and *An. coluzzii* from Burkina Faso available in Ag1000G was elucidated; of the 41,920 SNPs, 41,904 were present and thus taken forward to the next step. A final filtering step was introduced to focus on loci under selection in field populations. To do this, the top 5 % of the calculated H12 statistic for Burkina Faso populations included in Ag1000G were extracted for *An. gambiae* and *An. coluzzii* (Figure 5), and only those eQTL related SNPs present in the top 5 % H12 windows were considered. As previously reported, these sweep regions occur around *kdr, Rdl*, AGAP006222 (a UGT), the *CYP6P* cluster, the GSTE cluster and *CYP9K1* (Grau-Bove, et al. 2020; Clarkson, et al. 2021; Lucas, et al. 2024; Nagi 2025).

**Figure 4:**
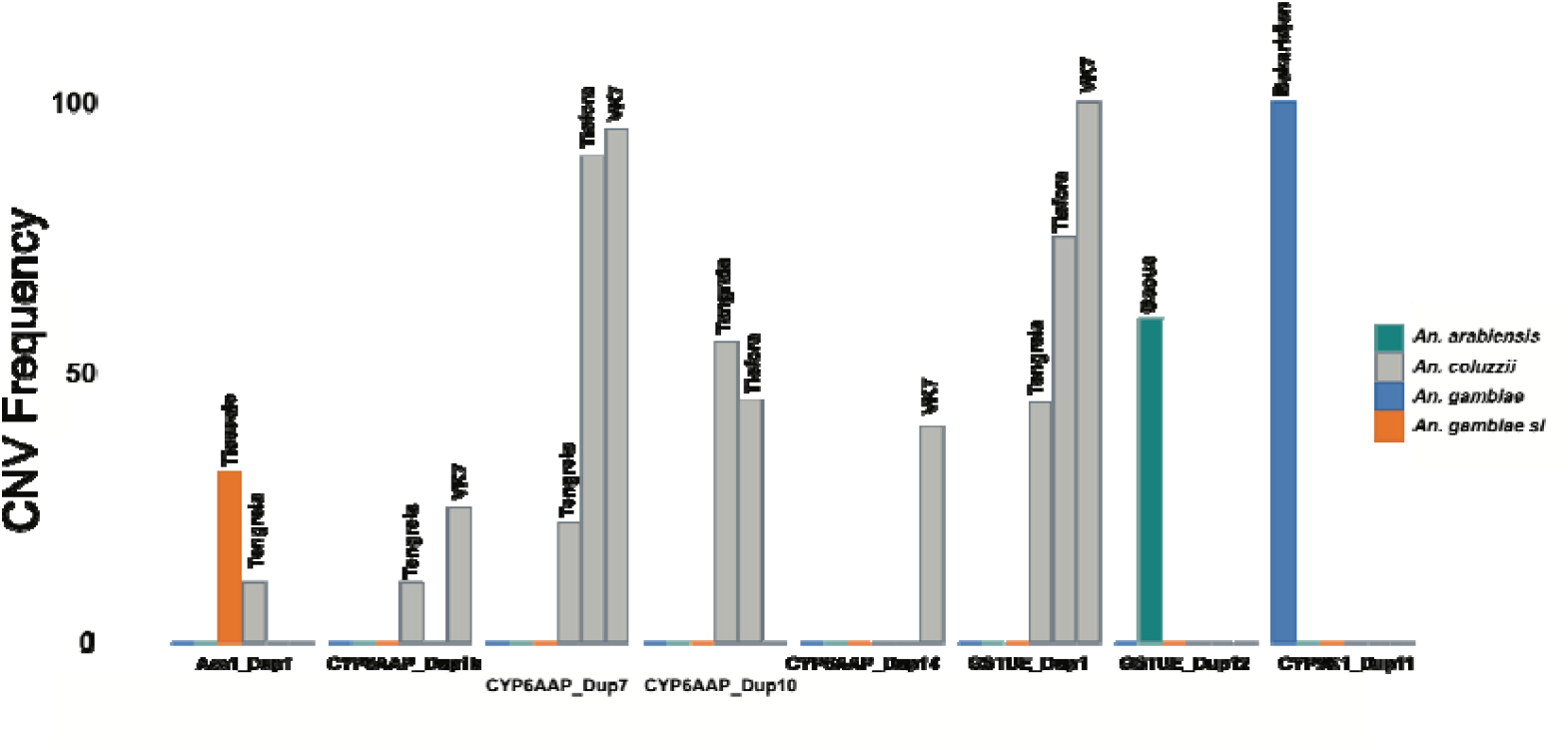
Copy number variations. CNV frequencies (y-axis) in each population (x-axis) of the major described duplications. Nomenclature as in [37]. *An. arabiensis* are shown in teal, *An. coluzzii* grey, *An. gambiae* blue and *An. gambiae* s.l. orange.

**Figure 5:**
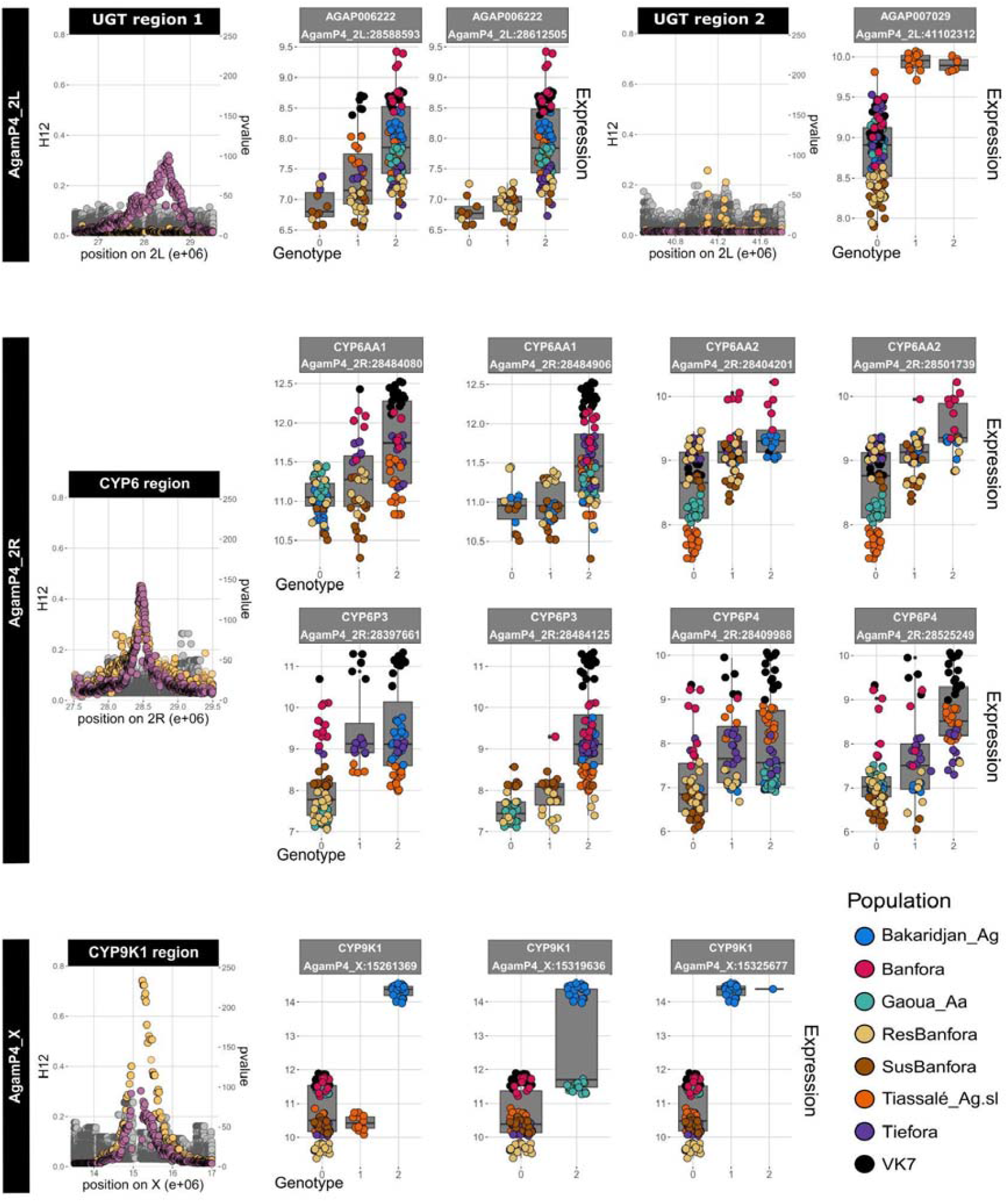
H12 and eQTL information for pyrethroid resistance-related transcripts. Dot plots showing Ag1000G H12 peak regions of interest for the two UGTs (AGAP006222 [UGT region 1], AGAP007029 [UGT region 2]), and the pyrethroid metabolisers *CYP6AA1, CYP6AA2, CYP6P3, CYP6P4* and *CYP9K1* (left y axis) on chromosomes 2L, 2R and X are shown with *An. gambiae* in pink and *An. coluzzii* in yellow along with eQTL p-values (right y axis) in grey. For each region of each chromosome, representative SNPs of interest showing increases in normalised read count (y axis) with genotype (x-axis) are shown as box plots with overlayed points. Genotype 0 = reference allele, 1 = heterozygote and 2 = derived. Points are coloured by population as shown in the key. SNP locations are given in the following format = AgamP4_chromosome:SNP locus.

After the above steps, a total of 1,008 eQTL-related SNPs were found within selective sweeps identified in the 2012 and 2014 Ag1000G samples from Burkina Faso: 771 in *An. coluzzii* and 621 in *An. gambiae*, with 384 of these SNPs being shared between the two species (Supplementary Table 8). Interestingly, 383 of the 384 shared SNPs are linked to expression of the *CYP6* cluster, with one linked to *ABCB1* expression. In total, the 1,008 SNPs relate to eQTLs associated with 34 resistance-related genes (Supplementary Table 7, Supplementary Table 8), including: *CYP9K1, CYP6AA1, CYP6P3* and *CYP6P4* all linked to pyrethroid resistance (Vontas, et al. 2018; Yunta, et al. 2019; Njoroge, et al. 2022) and two UGTs also linked with resistance (AGAP006222, AGAP007029) (Figure 5). eQTLs related to these transcripts were then explored further through plotting genotype against gene expression for each population (Figure 5, Supplementary Figure 11). eQTLs linked to *CYP9K1* show the highest gene expression in the *An. gambiae* Bakaridjan colony across three different SNPs. Most colonies show the wild type allele at all three loci, apart from the *An. gambiae* and *An. arabiensis* colonies. In the CYP6 region, several interesting associations can be seen. The derived allele of the first *CYP6P4* SNP for instance, is found across all three analysed species, but with the highest gene expression in *An. coluzzii. CYP6P4* has two SNP groups, with higher expression in VK7, Tiassalé_Ag.sl and Tiefora. *CYP6P3* has two associated SNP groups associated with lower expression in *An. arabiensis* and the susceptible Banfora and higher expression in *An. gambiae* and the *An. coluzzii* populations. Across the CYP6 region, the susceptible and resistant Banfora populations are relatively equally distributed across found alleles and show no differences in gene expression across the variants. However, interestingly, the SusBanfora and ResBanfora colonies (in which SusBanfora arose via the loss of the resistance phenotype in the original ResBanfora population) show evidence of association of *CYP6AA1* and *AA2* SNPs with the resistance phenotype: the derived form for 3 of the 4 SNPs in these genes is found at higher frequency in the resistant population and homozygotes for the derived form are absent in the susceptible strain. In general, the eQTL associations within this region are consistently characterised by a higher expression in the derived allele that is here mainly explained by VK7 and Banfora. The UGT AGAP006222 is associated with numerous SNPs, mostly driven by the susceptible *An. coluzzii* population, whilst AGAP007029 has two linked SNPs associated only with Tiassalé_Ag.sl.

## Discussion

*Anopheles* colonies are key resources in malaria research, being essential for studies on mosquito biology and also for screening and evaluating new control tools. Stable, fully genetically characterised insecticide resistant populations are rare but invaluable in linking results from laboratory and field experiments. Here, we build on previous transcriptomic and phenotypic studies on colonies of the *An. gambiae* species complex from Burkina Faso (Williams, et al. 2019; Williams, Ingham, et al. 2022) by incorporating whole-genome sequencing and using this to identify putative eQTLs. We provide evidence of potential introgression between species and generate data from further sampling of wild caught individuals to suggest that these shared haplotypes are not a result of laboratory contamination events but are likely occurring naturally by hybridisation within the species complex.

The evolutionary history of the *An. coluzzii* populations in this study appears to be independent of collection site or year of collection, with the colony Tiefora showing low relatedness to the other *An. coluzzii*. These data are in agreement with RNAseq and PBO-pyrethroid data showing that Tiefora displays lower reliance on cytochrome P450s than VK7, Banfora or Tiassalé_Ag.sl (Williams, et al. 2019; Williams, Ingham, et al. 2022) and thus, likely has differing insecticide resistance mechanisms. Furthermore, numerous cryptic species of *Anopheles* have been reported in Africa (Crawford, et al. 2016; Zhong, et al. 2020; Caputo, et al. 2024; Daron, et al. 2024) and indeed from the same region in Burkina Faso as the population origin (Tennessen, et al. 2021), reflecting a highly diverse species group, some of which is captured within the colonised populations. The diversity seen here underlines the necessity to test new vector control tool measures in multiple mosquito colonies and/or field populations to ensure variations in phenotypic responses are captured.

Inversions on chromosome 2 are a common feature of *Anopheles* populations (Love, et al. 2020), and have been linked with a variety of adaptive phenotypes such as resistance to desiccation [66] and susceptibility to *Plasmodium* infection (Riehle, et al. 2017). 2La is present in all populations at- or near-100 % frequency, whilst 2Rb and to a lesser extent 2Rc are at medium to high frequency in all populations except Tiassalé_Ag.sl; all three inversions have been significantly associated with resistance to pyrethroids (Ibrahim, et al. 2021; Ingham, Tennessen, et al. 2021; Ibrahim, et al. 2023). Copy number variations are evident across a number of previously described sites linked to pyrethroid resistance (Lucas, Miles, et al. 2019), including CYP9K1 in *An. gambiae* and CYP6AA1/AA2 in *An. coluzzii*. In addition to sites linked to resistance, well known immune genes including *TEP1* (Blandin, et al. 2004) and *APL1C* (Zmarlak, et al. 2024) show apparent duplications in this cluster, potentially indicating the strong selective pressure imposed by *Plasmodium* on these mosquitoes.

Evidence of selection is clear in the field-caught Tengrela (F) population, whilst bottle-necking in the colony populations appears to drive numerous instances of low-diversity regions. A sweep around the *CYP9K1* locus in Tengrela (F) reflects similar findings in multiple populations in the current release of Ag1000G (Anopheles gambiae Genomes 2020; Xue, et al. 2021). The H12-detected sweep found in the Tengrela (F) population centres on a synonymous SNP in *CYP9K1*, which could perhaps indicate changes to mRNA stability and translational efficacy (Hunt, et al. 2009) whilst the iHS statistics shows this peak to be upstream of *CYP9K1*. Similarly, selective sweeps in the *CYP6* locus are well described and attributed to the importance of this subfamily of P450s in detoxifying a variety of insecticides, including pyrethroids (Yunta, et al. 2016; Yunta, et al. 2019; Njoroge, et al. 2022). A signal around *Keap1* is observed in the colony populations, similar to what is seen in a longitudinal WGS from Burkina Faso (Kientega 2023). *Keap1* is a ubiquitin ligase that responds to external oxidative stress, allowing *cap’n’collar* to translocate to the nucleus and bind *Maf-S/Nrf* to trigger a stress response pathway (Ingham, et al. 2017). *Maf-S* has been shown to be up-regulated in multiple transcriptomic datasets, and when knocked down via RNAi increases susceptibility to pyrethroid insecticides likely through regulating metabolic resistance-related gene expression (Ingham, et al. 2017), indicating peaks around this loci could be directly linked with pyrethroid resistance. Although iHS and H12 identified peaks with resistance-associated alleles, the strength of the signal varied between populations, which could be due to extended reduced haplotype diversity through bottlenecking on colonisation and later colony contractions.

Introgression between the *An. gambiae* complex is well reported and hybrids are routinely seen at low frequency in field populations, indicating incomplete species divergence (Clarkson, et al. 2014; Fontaine, et al. 2015; Vicente, et al. 2017). Indeed, introgression has been shown at resistance-related loci including *kdr* (Clarkson, et al. 2014) and the *resistance to dieldrin* loci (Grau-Bove, et al. 2020). Here, multiple haplotypes appear to be shared between sister species, suggesting potential introgression. The I1527T-V402L haplotype, previously reported in *An. coluzzii* (Clarkson, et al. 2021; Kientega 2023) and *An. gambiae* (Collins, et al. 2019), is present in both the colony *An. arabiensis* and wild-caught *An. arabiensis* from the Cascades region in Burkina Faso with overlapping haplotypes to *An. coluzzii* colony populations from this region. Positive selection was detected around the *GSTE* locus, potentially driven by four non-synonymous SNPs in *GSTE1* and *GSTE7*, rather than mutations previously linked to pyrethroid resistance in *GSTE2* (Lucas, Rockett, et al. 2019; Diallo, et al. 2022). A mutation in *GSTE7*, L207I, is shown here to be significantly associated with deltamethrin survival, specifically when heterozygous. The N-terminal domain of GSTs is the xenobiotic hydrophobic binding domain (H-site), typically located between residues 106 and 220 (Oakley 2011), L207I falls directly in this region, and hence this substitution may impact insecticide or metabolite binding activity. Furthermore, *GSTE7* is included in several GST duplications (Gstue_Dup6-11) detected in Ag1000G (Lucas, Miles, et al. 2019), further linking this to field-based pyrethroid resistance. However, given the *GSTE7* and *GSTE1* SNPs are in complete linkage it is equally possible that SNPs in *GSTE1* are driving this association. In addition to resistance-related loci, *TEP1* appears to be shared between *An. arabiensis* and *An. coluzzii. TEP1* is a complement-like protein and has been shown to be a major factor involved in the mosquitos’ immune response to Plasmodium infection (Blandin, et al. 2004; Klug and Blandin 2023). *TEP1* is repeatedly found up-regulated in resistant populations (Ibrahim, et al. 2023; Ingham and Nagi 2024) and is the focus of a number of selective sweeps (Anopheles gambiae Genomes 2020; Tennessen, et al. 2021; Xue, et al. 2021) and haplotypes associated with resistance (Lucas, et al. 2023). Here, *TEP1* is shown to have no impact on pyrethroid resistance, indicating this sweep is likely driven by immune response to the malaria parasite and not insecticide exposure, in agreement with its described function (Blandin, et al. 2004).

The cis-eQTLs identified here may indicate putative regulatory loci associated with genes previously implicated in insecticide resistance, but these findings require further experimental validation. Neither the *GSTE* cluster, nor the *kdr* locus, are represented in these eQTLs, suggesting non-synonymous mutations might be driving pyrethroid-related resistance in line with previous expectations (Opondo, et al. 2016; Lucas, Miles, et al. 2019; Clarkson, et al. 2021; Diallo, et al. 2022). Similarly, no cis-eQTL loci were found associated with *CYP6M2*, whilst *CYP6P4* had clear cis-eQTL regions consistent with previously published data (Wagah, et al. 2021; Fotso-Toguem, et al. 2022). The *CYP9K1* eQTL is largely driven by *An. gambiae*, which is in line with this species exhibiting more CNV around this locus compared to *An. coluzzii* (Lucas, et al. 2023); however, RNAseq from multiple *An. gambiae* populations would be necessary to confirm this. The use of colony material limited the resolution of the current study precluding the identification of trans-eQTLs. Additionally, only biallelic SNPs were explored, limiting the ability to identify transposon insertions previously associated with insecticide resistance-related sweeps (Weedall, et al. 2020).

Overall, our findings demonstrate the value of colony populations for extrapolating laboratory results to field conditions, with several key observations validated in field-caught mosquitoes. We provide evidence of introgression at insecticide resistance associated loci across sister species, offering important insights into the spread of resistance and emphasising the need to evaluate new vector control products across multiple populations. These results reveal the porous species boundaries that facilitate adaptive gene flow amongst West African malaria vectors and highlights the critical importance of integrating genomic surveillance with vector control programmes to anticipate and mitigate the dissemination of resistance alleles. Finally, the high genetic diversity observed within laboratory colonies has important implications for the long-term efficacy and stability of gene drive strategies.

## Methods

### Mosquito samples and sequencing

Colonised insecticide resistant mosquitoes originally collected in the Cascades region of Burkina Faso or in southern Côte D’Ivoire were used for sequencing and are described in full by Williams et al (Williams, et al. 2019; Williams, Ingham, *et al*. 2022). Briefly, six resistant colonies and a field caught population were used in this study. Further, a previously published dataset from a susceptible and a resistant population deriving from the Banfora colony were included (Ingham, Tennessen, et al. 2021). The colony populations included are as follows: *An. gambiae* collected in 2015 (Bakaridjan, Burkina Faso); three *An. coluzzii* populations (Banfora (2015), Tiefora (2018) and VK7 (2014)) originating in Burkina Faso, one *An. gambiae* s.l. population from Côte D’Ivoire (Tiassalé_Ag.sl) collected in 2013 and one *An. arabiensis* collected in 2018 (Gaoua_Aa). The field colony here was collected in 2019 and is named Tengrela (F). The mosquitoes were reared at Liverpool School of Tropical Medicine (LSTM) under standard insectary conditions of 28°C, 80 % humidity with a 12:12 light:dark cycle and one-hour dawn:dusk, with the exception of Tengrela (F) which were reared in CNRFP, Burkina Faso. For this study, twenty individual females from each colony reared at LSTM were killed through freezing and total DNA extracted for each female using a DNAeasy Blood and Tissue kit (Qiagen) as previously described (Ingham, Tennessen, et al. 2021). Tengrela samples were stored on silica and shipped to LSTM where ten individual females had DNA extracted as described. The Banfora resistant and susceptible sequencing data used in this study was taken from a previous publication (Ingham, Tennessen, et al. 2021). Total extracted DNA was then quantified using Picogreen and submitted to the CeGaT sequencing centre (Tübingen, Germany) for 8GB flex sequencing with Illumina DNA library prep. In total 110 samples were sequenced within this project and 10 resistant Banfora and 20 susceptible and resistant taken from a prior publication (Ingham, Tennessen, et al. 2021), all sample fastq files can be found at the SRA under accessions PRJNA1149950 and PRJNA764501. The number of reads for sequencing generated within this project ranged from 152966234 to 83890472 (Supplementary Table 1).

### Genome alignment and quality control

The 160 fastq files were downloaded and multiple lane sequencing merged. The complete fastq files were then aligned to the *Anopheles gambiae* PEST assembly 4.13 from VectorBase (Giraldo-Calderon, et al. 2015) release 55 using BWA MEM v0.7.17 with default parameters (Li and Durbin 2010). The percentage of mapped reads ranged from 76.35 % to 99.29 %, with a mean of 97.08 % (Supplementary Table 1). Duplicates were then marked and the bam file reordered using Picard 2.9.0 (http://broadinstitute.github.io/picard/) with default parameters and indexed using samtools v1.9 (Li, et al. 2009). Variants were called using the best practice GATK pipeline with HaplotypeCaller v4.1.6 (McKenna, et al. 2010). BCFtools v1.9 (Li 2011) was then used to remove calls with missingness >0.05, minor allele frequency of > 0.01 and depth and quality scores > 5 and 20 respectively. PCA analysis was performed using plink2 (Purcell, et al. 2007) and visualised using ggplot2 (Villanueva 2019) in R. Three samples (QJIUK036A3, QJIUK169AK and QJIUK068A6) out grouped from expected in the PCA and were removed from further analysis, resulting in 157 samples within the final variant call file (vcf). For tools requiring haplotypes, the vcf was then phased using Beagle v5 (Browning, et al. 2021) and maps produced as part of Ag1000G. A biallelic vcf was produced using BCFtools v1.9 -M 2 flag.

### SNP effects

To determine the impact of the identified SNPs, two approaches were taken. Firstly, SNPEff (Cingolani, et al. 2012) was used to annotate SNPs across the entirety of the vcf. The second approach used the Ag10000G data (Anopheles gambiae Genomes, et al. 2017), with in-country allele frequency and SNP-directed modifications extracted.

### Phylogenetic analysis

To remove confounding variables such as inversions, chromosome 3L was used for all phylogenetic analysis. Two methods were used to determine the phylogenetic relationships of the samples. In each instance the vcf file was converted to fasta format using the vcf2phylip python script retaining the major allele. Firstly, SplitsTree (Huson and Bryant 2006) was used to create an unrooted phylogenetic network with 10,000 bootstraps. Maximum likelihood trees were also constructed using RaXML v8.2.4 (Stamatakis 2014) using the GTRGAMMA model.

### Admixture analysis

Admixture software (Alexander, et al. 2009) v1.3 was used to infer population ancestry amongst the mosquito populations. To determine the correct cluster number, k 1 to 10 were tested. To avoid error in the parameterisation, 1,000 bootstraps were used. The optimal cluster number was determined based upon cross validation error and was determined to be k = 6 or k = 7.

### Inversion status

Inversions were detected using previously published tagging SNPs for the inversion (Love, et al. 2020). In many instances not all tagging SNPs displayed the same pattern or hetero/homo-zygosity and therefore, incidences where >80 % of tagging SNPs followed the same pattern was classified as an inversion.

### CNV scan

CNV scan was done as previously described for all populations (Lucas, Miles, et al. 2019).

### Inferring selective sweeps

Patterns of selective sweeps were analysed using a combination of iHH12 and iHS statistics (Voight, et al. 2006; Garud, et al. 2015). The iHS and iHH12 statistics were calculated using selscan v2.0 (Szpiech 2024) with default parameters to infer regions of extended haplotype diversity on each individual chromosome. Before each, the vcf was split into individual populations and low frequency variants removed following selscan definition. The top 1 % of the absolute iHS and standardised iHH12 statistics were considered as the final candidate sets. iHH12 was standardised by splitting alleles into 1 % bins and performing the scale function within bins using the dplyr package (Wickham H 2025).

### Diversity statistics

F_ST_, nucleotide diversity and TajimasD were calculated using vcftools v0.1.17 (Danecek, et al. 2011). For each population combination, both per-site and windowed values (10 kb windows with 5 kb steps) were calculated. F_ST_ was calculated using the -weird-fst-pop function in vcftools, comparing all populations in a pairwise manner.

### Introgression

To identify potential areas of introgression the F_ST_ and ABBA-BABA statistics (Durand, et al. 2011) in sliding windows were used. ABBA-BABA’s Patterson’s D and f_4_-statistics used in population genetics allow the relationship between four taxa to be determined. In a phylogenetic relationship following (((P1,P2),P3),O) a positive D statistic infers introgression between P2 and P3, whilst a negative D statistic infers introgression between P1 and P3. The outgroup used in this analysis is the Angola population from Ag1000G (Anopheles gambiae Genomes, et al. 2017), being geographically and temporally distant from the populations used in this study. For this study, the python script ABBABABAwindows.py from Martin et al. was used (Martin, et al. 2015), with 100 kb non-overlapping windows and the f_dM_ statistic used as recommended.

To identify putative regions of introgression, genomic windows falling within the upper 5% of absolute |fdM| values and the lower 5% of F_ST_ values were identified, and their intersections taken as candidate introgressed regions. fdM, derived from ABBA–BABA statistics, highlights genomic regions deviating from the expected phylogenetic relationship among populations, whereas F_ST_ quantifies genetic differentiation between population pairs. To minimise bias arising from variation in the number of informative sites across the genome, windows were filtered to retain only those exceeding minimum data thresholds (ABBA–BABA: sites ≥ 10 and F_ST_: N_VARIANTS ≥ 10 where available). In addition, F_ST_ outliers were defined using empirical, data-availability–corrected thresholds: for each focal– comparator population pair, windows were stratified into deciles based on a proxy for information content (N_VARIANTS per window), and the lower 5th percentile of F_ST_ was computed within each stratum. Windows were then classified as “low-F_ST_” if their F_ST_ was at or below the stratum-specific threshold, ensuring that low-F_ST_ calls were not driven by windows with reduced SNP density or poor coverage. Thresholds were calculated only on autosomes. As the combined threshold approach assumes that fdM and F_ST_ provide largely independent information, this assumption was explicitly evaluated. First, the genome-wide association between fdM and F_ST_ was quantified for each focal population using both Pearson’s product–moment correlation and Spearman’s rank correlation. Second, to test whether fdM outlier windows were preferentially associated with reduced genetic differentiation between the recipient population and the inferred donor, a permutation test was performed. For each focal population, fdM outlier labels were randomly reassigned within population-pair and chromosome strata, while genomic window positions and F_ST_ values were held fixed. For each permutation, the number of fdM outlier windows overlapping regions of donor-specific low F_ST_ was recorded, generating a null distribution that accounts for background relatedness and genomic heterogeneity. The observed number of donor-supported fdM outliers was compared to this null distribution to assess enrichment. Together, these analyses support the combined use of fdM and diversity-corrected F_ST_ thresholds as a conservative criterion for identifying candidate introgressed regions (Supplementary Table 3).

### Haplotype maps and clustering

In areas of introgression, haplotype maps were used to explore the relationship in the identified windows of introgression and/or selective sweeps. To visualise these regions a custom script was written in R using ape (Paradis and Schliep 2019) and pegas (Paradis 2010) packages. Briefly, the region was extracted from the vcf file using vcftools and read into R using read.vcfR. The resultant file was then converted to a fasta file using vcfR2DNAbin and haplotype data produced following instructions in the pegas package (Paradis 2010). For cluster visualisation R (Team 2021) was used as follows: Genotypes were parsed to extract the GT field and split into two alleles per diploid sample, generating one row per haplotype. SNPs that were invariant or entirely missing were excluded. Missing values were imputed using SNP-wise means, and data were scaled prior to clustering. Pairwise Euclidean distances were calculated and hierarchical clustering was performed using Ward’s minimum variance method with the stats package. Cluster membership was inferred using silhouette optimisation (cluster package) (Bhat K S 2025) and adaptive dynamic tree cutting (dynamicTreeCut package) (Langfelder, et al. 2008). Population annotations were inferred from sample identifiers. Haplotype heatmaps were generated using pheatmap (Kolde 2025), and dendrograms with cluster-coloured branches and population-coloured tips were visualised using ggplot2 (Villanueva 2019) and ggdendro (de Vries 2025).

### Enrichment Analysis

Enrichment analysis was performed on VectorBase (Giraldo-Calderon, et al. 2015) for GO terms and KEGG. The cut-off for significance was set at p <= 0.05 after Bonferroni correction.

### Mosquito rearing for validation

Tiassalé_Ag.sl and Tiefora mosquitoes were reared at a constant temperature of 27 ± 2 °C and 75 ± 10 % humidity, under a 12:12 light:dark cycle with one hour dawn:dusk at Heidelberg University Hospital. Mosquitoes were maintained in milliQ water (RephiLE PURIST) from eggs to pupae stage. Larvae were supplemented with cat and fish food, with adult mosquitoes then fed on 10 % sucrose sugar (w/v) in milliQ water, if not indicated otherwise. For all experiments, presumed mated female mosquitoes were used at 3-5 days old unless otherwise stated.

### RNA extractions and cDNA synthesis

For all work involving RNA, five to seven whole, adult female mosquitos were snap frozen, pooled and homogenised. Their RNA was extracted (Arcturus™ PicoPure™ RNA Isolation kit) according to the manufacturer’s instructions, all samples were treated with DNase I (RNase-Free DNase Set) during this clean-up to remove residual DNA. For mRNA production, RNA (4⍰µg) from each biological replicate was reverse transcribed using Oligo dT (Invitrogen) and Superscript III (Invitrogen) according to manufacturer’s instructions.

### dsRNA synthesis and microinjection

dsRNA was generated as previously described (Ingham, et al. 2018). Briefly, Tiassalé_Ag.sl cDNA from three to five-day old females was used as a template. dsRNAs of target genes were generated using Phusion polymerase (Thermo Scientific). All reactions were conducted with the same PCR cycle conditions: 98°C 30s, 35 cycles of (98°C for 10s, annealing temperature for 10 s, 72 °C for 1m), 72°C 45s. All primers used for dsRNA generation contained a bacterial T7 sequence (5’ – 3’: TAATACGACTCACTATAG) at their 5’ end to allow for dsRNA generation. GFP: Fwd: AGAACGTAAACGGCCACAAGTTC, Rev: AGACTTGTACAGCTCGTCCATGCC. TEP1: Fwd: CTGAAAGCGCTGACCAA, Rev: TATGTAGCTGGCACAGACC. Reactions were purified using a QIAquick® PCR Purification kit. A total of 1 - 1.5 µg purified DNA of a single band was used as template for dsRNA generation with a commercial kit (MEGAscript™ T7 kit). Two reactions were set up for one target and pooled to increase yield, remaining steps were performed according to the manufacturer’s instructions. Following purification of generated dsRNA (MEGAclear™ Kit), the templates were adjusted to a concentration of 3 µg/µl by vacuum evaporation (CentriVap®). dsRNAs were stored at -20 °C until further use. Subsequent injections were carried out at a 69nL volume through the thorax using a NanoInject II.

### qPCR

The primer efficiencies of the qPCR primers (Table 2) were initially verified to be between 80 and 120 % using Tiassalé_Ag.sl cDNA of three-to-five-day old females as input and serial dilutions of 1:5. To quantify the transcription levels of desired targets, three biological replicates of cDNA, with three technical triplicates each were analysed per group. As master-mix, the SYBR Green qPCR (Thermo Scientific) mix was used according to the manufacturer’s instructions with 2 ng total cDNA as input. The genes 40S ribosomal protein S7 (S7) and elongation factor 1-alpha (EF) were used as internal references for normalizing the target transcript levels. Following 40 amplification cycles, a melt-curve of the amplified products from 55 to 90 °C was recorded in 0.5 °C steps to rule out influences of off-target amplifications. Ct values were calculated using the delta delta Ct method. All qPCR was carried out with the following conditions: 3⍰min at 95⍰°C, with 40 cycles of 10⍰s at 95⍰°C and 10⍰s at 60⍰°C.

**Table 2:**
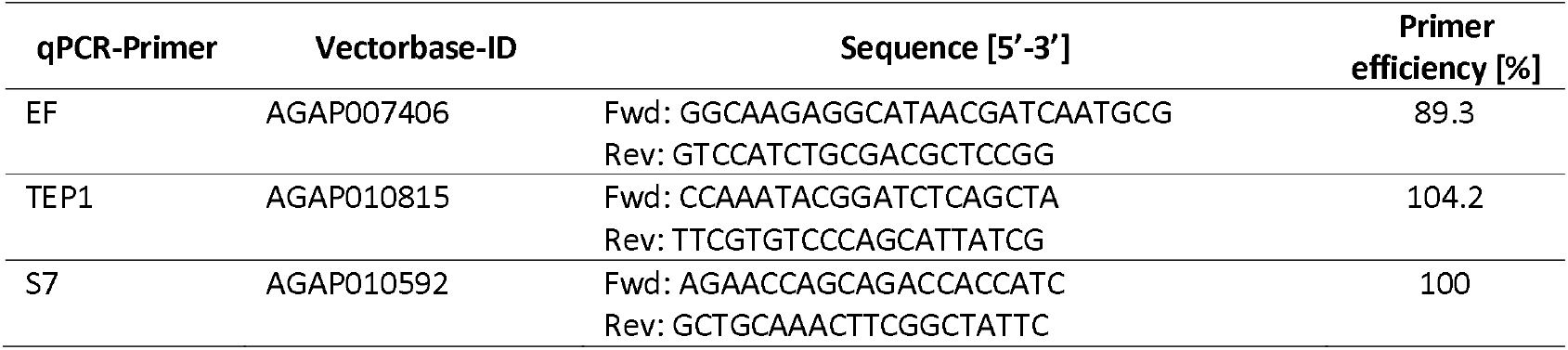
qPCR-primers used for quantification of respective transcripts using SYBR-green.

### Protein Quantification

Five whole, adult Tiassalé_Ag.sl mosquitoes were pooled per replicate and ground in 50 µl freshly prepared protein extraction buffer (50 mM Tris, pH 8.0; 1 % NP40; 0.25 % sodium deoxycholate; 150 mM NaCl; 1 mM EDTA; 1X protease inhibitor mixture (Roche, Basel, Switzerland)). Samples were homogenised for ten intervals at 60 % force using a sonicator and incubated on ice for 30 min. Samples were centrifuged at 21.100 x g for 10 min, and supernatant transferred into a new tube. The concentration of proteins was determined using a Coomassie solution and comparison to a bovine serum albumin (BSA)/PBS standard series, as described in the Bradford assay for quantification of proteins. 13 µl of respective protein solution, 5 µl 4 x Laemmli buffer and 2 µl 1 M DTT were mixed and incubated at 70 °C for 10 min. The whole volume was loaded onto 4 – 20 % Tris-Glycine gels and ran in SDS-buffer at 100 V for 40 - 50 min, followed by semi-dry blotting onto a 0.2 µm nitrocellulose membrane for 7 min at 1.3 A using the mixed molecular weight program. Membranes were briefly rinsed with PBS-T (PBS + 0.1 % (v/v) Tween20) and stained with Ponceau solution for 5 sec to verify the presence of proteins. Membranes were repeatedly rinsed with tap-water to remove Ponceau staining and incubated for 1h at RT in blocking solution (5 % (w/v) milk powder in PBS-T), followed by incubation with one or more primary antibodies over night at 4 °C on a rotator. Between each antibody, membranes were washed 3x for 5 min in PBS-T at RT on a shaker. Ultimately membranes were incubated with secondary antibodies for 1-2 h in the dark, at RT on a rotator, washed 3 x 5 min with PBS-T, 1 x 15 min with PBS-T and 1 x 5 min with PBS and imaged on a Li-Cor Odyssey. Respective bands were quantified using the built-in analysis function in ImageStudio. The levels of target proteins were normalized to levels of the housekeeping gene α-tubulin.

### Resistance Testing

To determine the resistance status of mosquitos, 25 healthy females were aspirated into tubes for a standardized WHO-tube test at diagnostic dose alongside a control, as previously described (WHO 1998).

### Longevity

Females were stored in cups with no more than 20 females per cup and mortality scored daily.

### gDNA extraction

gDNA was extracted from whole mosquitoes or individual legs through homogenising and boiling for 30mins at 95°C in STE buffer.

### LNA validation of I1527T

To validate the presence of I1527T, a previously published LNA diagnostic was used [15]. Briefly, reactions were set up using 1⍰× ⍰Luna Universal qPCR Master Mix (NEB), 0.1⍰μM for each probe (IDT), 0.2⍰μM of primers and 1–2⍰μL of DNA extract in a total reaction volume of 10⍰μL. Reactions were run on an AriaMX qPCR cycler with the following conditions: 95⍰°C for 3⍰min, followed by 40⍰cycles of 95⍰°C for 5⍰s and 60⍰°C for 30⍰s. FAM probes correspond to 1527T, whilst HEX corresponds to 1527I. To validate the presence of I1527T in the *An. arabiensis* colony, Gaoua_Aa, historic DNA samples collected over multiple generations since colonisation were explored at LSTM from generation 8 to generation 54 (G8, G17, G26, G31, G38, G46, G54). To confirm whether this mutation was detected within Burkina Faso, 95 field collected samples previously identified as *An. arabiensis* via SINE PCR were shipped from CNRFP, Burkina Faso. The collections were carried out in Tiefora village (10.624116669/-4.55335) between July and September 2021, one individual from Tengrela (F) in September 2021 (10.65188333/-4.82441667) and in Ouagadougou at two collection sites, Zogona (12.3786/-1.48752) and Karpala (12.31929/-1.50171) in November and September 2022 respectively.

### Sanger sequencing of *kdr*

To verify the positive results from the field caught *An. arabiensis*, primers were designed to span the region around I1527T. Briefly, PCR reactions for the amplification of the target site were set up as follows: 0.25 μl of 5 U/μl DreamTaq DNA Polymerase (ThermoFisher Scientific), 2.5 µl 10X DreamTaq Buffer (ThermoFisher Scientific), 0.5 µl 10mM dNTPs, 1.5 µl of 10 µM primers with 1 µl of the sample DNA in a reaction volume of 25 µl. Reactions were run on a Biometra TOne PCR thermocycler at 98 °C for 30 s, followed by 30 cycles of 98°C for 10 s, 55°C for 30 s and 72°C for 40 s; this was followed by a final extension step at 72°C for 5 min. Primer sequences were as follows: forward: ACCGGATGTAAATGCTTGCA, reverse: CCGCCACTGGAAAGAATGGA. Resulting amplicons were purified using the QIAquick PCR purification kit (Qiagen, Germany). Sanger sequencing was done using the PCR primers by GENEWIZ (Leipzig, Germany).

### GSTE haplotype determination

Validation of L207I mutation circulating in field populations of *An. arabiensis* was done in the same field samples used for the *kdr* mutation I1527T. For this, a partial fragment of 903 bp from the GSTE7 gene was amplified in the samples using the primers GSTE7_WS_F and GSTE7_WS_R (see Table 3). PCR reactions contained 1X Phusion HF Master Mix (Thermo Scientific), 0.5 μM of each primer and 2 μL of gDNA in a total volume of 50 μL. Amplification conditions included an initial denaturation at 98°C for 30 s, followed by 35 cycles of 98°C for 10 s, 63°C for 30 s, and 72°C for 15 s, with a final extension step of 10 min at 72°C. Samples showing a specific band were sent to GENEWIZ (Leipzig, Germany) for sequencing using the forward PCR primer. Electropherograms were visualized and analysed using FinchTV (Geospiza, Inc) and L207I mutations identified in the sequences.

To assess the association between the mutation L207I on the GSTE7 and pyrethroid resistance on Tiefora and Tiassalé_Ag.sl colonies, an allelic-specific PCR was developed and then applied to genotype this mutation on dead and alive mosquitoes exposed to insecticides. Initially a sub-sample of females from the Tiefora colony was used to isolate genomic DNA using 1X STE buffer. A partial fragment spanning the L207I mutation was amplified in the samples using the same primers for *An. arabiensis* (GSTE7_WS_F and GSTE7_WS_R) following the same PCR reaction mix and amplification conditions mentioned above. Samples showing a specific band were sent to GENEWIZ (Leipzig, Germany) for sequencing using both PCR primers and electropherograms were visualized and analyzed using FinchTV (Geospiza, Inc). Samples showing a clear peak for the genotype L/L, L/I and I/I were used as positive controls to validate an allelic discrimination PCR assay. Then, primers for an allelic discrimination assay were designed with mismatches to improve the specificity of the wild and mutant allele [60]. For this, a partial fragment of 279 bp was amplified for the wild type allele using the primers L207I_wt_F and L207I_R whereas mutant allele was amplified using a second forward primer L207I_mt_F (Table 3). Thus, for each sample, a separate PCR reaction was prepared for each allele. Each PCR reaction contained 1X Buffer Dream Taq (Thermo Scientific), 0.2 mM of dNTPs mix, 0.6 μM of each primer, 1.25 units of Dream Taq Polymerase, and 2 μL of gDNA in a total volume of 25 μL. The amplification conditions included an initial denaturation at 95°C for 5 min, followed by 40 cycles of 94°C for 30 s, 68°C for 30 s, and 72°C for 1 min, with a final extension step of 10 min at 72°C. A sample was classified as wildtype when this showed positive amplification only in the wt PCR, mutant when this showed positive amplification only in the mt PCR, and heterozygous when showed amplification in both wt and mt PCR reactions. For the insecticide exposures, healthy females from Tiefora and Tiassalé_Ag.sl were aspirated into tubes for a standardized WHO-tube test at 1X and 5X DD of deltamethrin, permethrin, and alpha-cypermethrin alongside a control, as previously described [97]. After 24 h, mortality was registered, and alive and dead mosquitoes were sorted and stored at -20°C. Genomic DNA was isolated for each mosquito using the 1X STE buffer and each sample was genotyped using the allelic-specific PCR developed for the mutation L207I on *GSTE7*. The frequency for the different genotypes was calculated in the alive and dead mosquitoes per colony and per insecticide. The frequencies observed were compared among them using the Fisher Exact Test and a p-value of 0.05 was used as threshold for significance (GraphPad Prism).

**Table 3:**
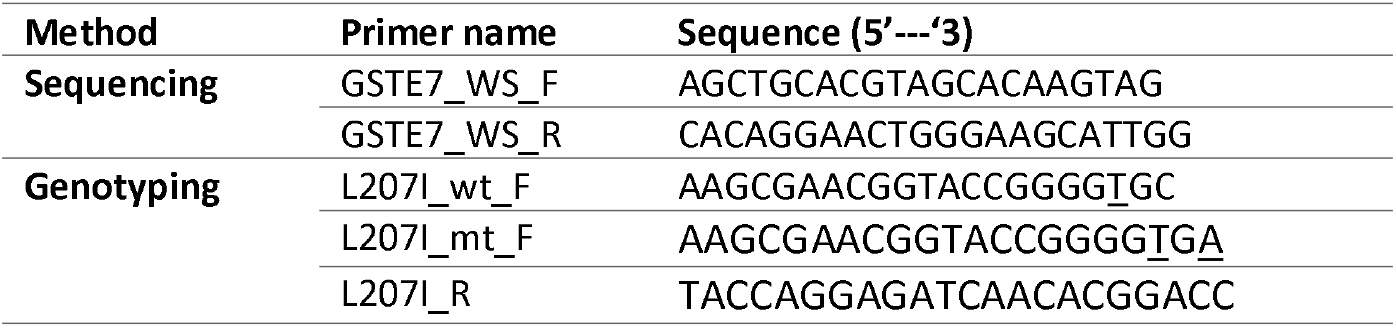
Primers for sequencing and genotyping of the mutation L207I on *GSTE7* from *An. coluzzii* and *An. gambiae* s.l. Mismatches including in the genotyping primers are highlighted with underlines.

### RNAseq of Tiassalé_Ag.sl colony

3–5-day old female Tiassalé_Ag.sl mosquitoes were killed by snap freezing and total RNA extracted from four pools of 7 mosquitoes using a PicoPure RNA Extraction Kit with DNAse treatment. RNA quantity and quality was checked using a NanoDrop and Bioanalyzer, respectively. Total RNA was then submitted to Eurofins (Ebersberg, Germany) for sequencing after polyA enrichment.

### eQTL analysis

For the eQTL analysis, RNAseq data for Banfora, BanforaSus, BanforaRes, Tiefora, Gaoua_Aa, VK7 and Bakaridjan_Ag were available from previous studies (Ingham, Tennessen, et al. 2021; Williams, Ingham, et al. 2022), whilst Tiassalé_Ag.sl was generated here. The fastq files were downloaded, aligned using Hisat2 v2.2.1 (Kim, et al. 2015) to the *Anopheles gambiae* PEST assembly 4.13 from VectorBase release 55 with default parameters. featureCounts from the subread package v2.0.3 (Liao, et al. 2014) was then used to produce a count file for all populations. As the transcriptomic data stems from different individuals than those used for the WGS, preventing direct correlations at the individual level, counts were inferred using a normal distribution constrained by the range of counts seen for each gene within that population. This approach keeps the overall distribution of expression but breaks any true genotype-expression linkage, ensuring that the observed effects are not based on inherited noise stemming from mismatched data, allowing for individual-level correlations during eQTL analysis. The generated counts file was then corrected for batch effects and library size using the limma v3.58.1 (Ritchie, et al. 2015) and DEseq2 v1.42 (Love, et al. 2014) packages respectively. eQTL analysis was then performed to identify cis-eQTLs only following standard practice as outlined in the MatrixEQTL package (Shabalin 2012) v2.3 in R. MatrixEQTL uses linear regression to identify associations between genotypes and gene expression levels at the individual level. The p--value cut off was defined as 0.05/total number of SNPs, hence p < 5.37e-9. The distance for a valid cis-eQTL was defined as 10,000 bp taken from a *Drosophila* paper showing that the majority of cis-eQTLs are found within this distance (Yang, et al. 2022). To determine an FDR cut-off, a permutation (n=100) test was carried out that scrambled the count data and hence the genotype:count relationship as previously described (Quigley, et al. 2009). The calculated p-values for each SNP were then passed to the qvalue package v2.34 and an FDR cut-off of 0.1 applied (Storey and Tibshirani 2003).

eQTLs were then extracted in the regions of known *a priori* insecticide resistance candidates, focusing on metabolic detoxification enzymes, chemosensory proteins, hexamerins, neuronal targets, alpha-crystallins and D7 salivary gland proteins (Supplementary Table 6). Families were extracted from VectorBase using PFAM IDs as follows: PF00067, PF00201, PF00135, PF00005, PF12848, PF03392, PF00011, PF03722,

PF00372 and PF01395. For eQTLs located in or around genes known to be implicated in insecticide resistance, the Ag3 dataset from the MalariaGEN project (v11.0.0) was used to extract allele counts across all sample sets originating from Burkina Faso (AG1000G-BF-A, AG1000G-BF-B, AG1000G-BF-C) using the ag3.snp_allele_counts function. In a second step, we extracted H12 statistics from the same sample sets to gather information whether eQTLs are located in regions of selective sweeps. This was done separately for *An. gambiae* and *An. coluzzii* but included all available cohorts of those two species (*An. coluzzii*: BF-09_Houet_colu_2014, BF-09_Houet_colu_2012; *An. gambiae*: BF-09_Houet_gamb_2014, BF-09_Houet_gamb_2012). For each chromosome, window size was calibrated with the ag3.plot_h12_calibration function and H12 statistics were then extracted with the ag3.h12_gwss for the specific window size. H12 outlier windows were determined as those belonging to the 95 %-quantile of the observed distribution. We displayed H12 distributions across the chromosomes using ag3.plot_h12_gwss and visually determined regions with naturally elevated H12 towards the telomeric regions that were then excluded from further analysis. Where SNPs were present as a linked block, only one SNP was chosen for visualisation.

---

**Supplementary Figure 1: Descriptive statistics for each population. A**. Nucleotide diversity and **B**. Tajimas D statistic for each population sequenced. The respective statistics are shown on the y axis and the x axis shows individual chromosomes. Each box plot represents a separate population, coloured as in the key.

**Supplementary Figure 2: Absolute iHS statistic for the colony populations**. iHS statistic (y axis) for the length of each chromosome (x axis) as indicated at the top of each panel. Colonies are coloured as previously, and the title of the graph indicates the colony. Displayed are the top 5 % of the statistic only, cut-off indicated below. Negative iHS are shown in black. Labels correspond to putative insecticide-resistant associated transcripts in regions of elevated iHS.

**Supplementary Figure 3: Absolute H12 statistic for all populations**. H12 statistic (y axis) for the length of each chromosome (x axis) as indicated at the top of each panel. Populations are coloured as previously, and the title of the graph indicates the population. Displayed are the top 5 % of the statistic only, cut-off indicated below. Labels correspond to putative insecticide-resistant associated transcripts in regions of elevated iHS.

**Supplementary Figure 4: iHS sweeps in the Tengrela (F) population centred on genes of interest**. Absolute iHS statistic (y axis) for each resistance-related sweep location in the field-caught Tengrela (F) population. The loci of the genes of interest were taken as the centre point +/-2Mbp (x axis). The dotted line represents the gene of interest: **A**. *kdr*, **B**. *CYP6P3*, **C**. *CYP9K1* and **D**. *GSTE2*. The yellow line represents positive iHS and black negative. Negative values signify selection on derived alleles while positive values are associated with selection on ancestral alleles.

**Supplementary Figure 5: H12 sweeps in the Tengrela (F) population centred on genes of interest**. Absolute H12 statistic (y axis) for each resistance-related sweep location in the field-caught Tengrela (F) population. The loci of the genes of interest were taken as the centre point +/-2Mbp (x axis). The dotted line represents the gene of interest: *GSTE2* and *CYP9K1*. The yellow line represents H12.

**Supplementary Figure 6: X chromosome ancestry informative markers for each population**. Ancestry informative markers used in Ag1000G were extracted from the vcf file and assigned the expected species. **A**. Bakaridjan_Ag, **B**. Banfora, **C**. Gaoua_Aa, **D**. Tengrela (F), **E**. Tiassalé_Ag.sl, **F**. Tiefora and **G**. VK7. Blue shows *An. gambiae* markers, pink shows *An. coluzzii*, orange shows *An. arabiensis* and green shows heterozygotes.

**Supplementary Figure 7: Summary of shared haplotype regions**. The number of shared haplotype regions in the top 5 % of windowed ABBA-BABA and bottom 5% of F_ST_ statistics across all populations. The intersection size (y) shows the number of regions shared between the populations listed below the x-axis. Each dot connected by a solid line shows the populations with the shared haplotypes. The bar chart to the left shows the total number of putative haplotypes shared for each population. *An. coluzzii* are purple, *An. gambiae* orange and *An. arabiensis* teal.

**Supplementary Figure 8: Dendrogram for loci with shared haplotypes**. Dendrograms for **A**. *CYP9K1*, **B**. *CYP6P* cluster, **C**. *CYP4G16*, **D**. *CSP* cluster, **E**. *Rdl* and **F**. *TEP1* branch length represents numbers of SNP differences. Clusters defined by optimal k as determined by a silhouette plot are represented through coloured branches and nodes labelled by species colours. For each, a heatmap showing wt (blue) and derived (green) alleles are shown to give an indication of the diversity within and between clusters. Colours at the side of the heatmaps represent the branch colours.

**Supplementary Figure 9: Mortality of Tiefora and Tiassalé_Ag.sl mosquitoes and association testing. A**. Mortality of Tiefora and **B**. Tiassalé_Ag.sl to WHO tubes of 1X and 5X DD of deltamethrin, permethrin and alpha-cypermethrin. **C-F**. Association study of L207I with resistance to 5X DD using WHO tube tests of three pyrethroids in the **C/D**. Tiefora for **C**. permethrin and **D**. alpha-cypermethrin and **E/F**. Tiassalé_Ag.sl for **E**. permethrin and **F**. alpha-cypermethrin. n shows number of mosquitoes screened, wt = wild-type, H = heterozygous, mt = mutant, no significance, as calculated by a Fisher Exact Test.

**Supplementary Figure 10: Knockdown of *TEP1* has no impact on pyrethroid resistance in Tiassalé_Ag.sl. A**. Bar graphs depicting transcript levels (y-axis) of *TEP1* after dsRNA injection (grey bars), normalized to the GFP-injected control (black bars) 72- and 96-h post injection. Depicted values are the average of three biological replicates with three technical replicates each, ± SD. Statistical comparison between control and RNAi group were calculated with unpaired t-test: * p < 0.05 **B**. Bar chart showing comparison of quantified *TEP1* (processed) levels between control and *TEP1* dsRNA injected mosquitos, after normalization of respective signal to the α-tubulin loading control. Western blot of TEP1 proteins in control dsGFP (left) and dsTEP1 (right) injected mosquitos over a time course, below is the α-tubulin loading control. d3 and d5 refer to day 3- and 5-post-injection. **C**. Kaplan-Meier-plot comparing the survival of non-injected (black) versus dsTEP1-injected mosquitoes (red). The vertical line represents time of injection. 50 female mosquitos were used per group. **D**. Observed 24 h mortality of pyrethroid-resistant after 1 h exposure to a control tube or 0.05 % deltamethrin in a WHO-tube assay. Percentage mortality (y axis) for control exposed for each injection group (black), dsGFP (grey) and dsTEP1 (red). Statistical comparison between control and deltamethrin exposed group were calculated with unpaired t-test. Sample size is indicated below bars. All experiments done with Tiassalé_Ag.sl.

**Supplementary Figure 11: H12 and eQTL information for pyrethroid resistance-related transcripts**. Dot plots showing Ag1000G H12 peak regions of interest for insecticide resistance-related transcripts (left y axis) across chromosomes are shown with *An. gambiae* in pink and *An. coluzzii* in yellow along with eQTL p-values (right y axis) in grey. For each region of each chromosome, representative SNPs of interest showing increases in normalised read count (y axis) with genotype (x-axis) are shown as box plots with overlayed points. Genotype 0 represents reference allele, 1 heterozygote and 2 derived. Points are coloured by population as shown in the key. SNP locations are given in the following format AgamP4_chromosome:SNP locus.

## Data Availability

WGS data has been deposited in SRA under accessions PRJNA1149950 and PRJNA764501 and RNAseq data under PRJNA1185152, accession numbers SRR31314079-82. All scripts are available in github: https://github.com/jubiology/eQTL-H12-analysis and https://github.com/VictoriaIngham/BurkinaWGS.

## Acknowledgements

This study was funded by the Deutsches Zentrum für Infektionsforschung (DZIF, TTU03.705), ERC Starting Grant (ReMVeC, Project Number 101075634), the Deutsche Forschungsgemeinschaft (DFG, German Research Foundation) – project number 240245660-SFB 1129 awarded to VAI and a DAAD fellowship awarded to JCL (91828456). Thanks to Wolfgang Huber, EMBL for computational infrastructure, Stephanie Blandin, CNRS/Inserm for the *TEP1* antibody and to Alistair Miles, University of Oxford for assistance with integrating Ag1000G data. We also thank Timo Federolf for assistance with the GSTE genotyping. We acknowledge CNRFP and IRSS for assistance with original colonisation of mosquito populations used in this study and LITE for mosquito rearing.

## Contributions

VAI conceptualised the study, performed all data analyses for the whole genome sequencing and drafted the manuscript. MM and PH prepared DNA for sequencing. JCL, ALB, MM and JH performed the lab validation of the *GSTE* and *kdr* haplotypes. JCL performed all Tiefora experiments. JH integrated eQTL data with Ag1000G data. ALB generated Tiassalé_Ag.sl RNAseq. JKK, JCL and ALB performed TEP1 experiments. AS and MG contributed field collected mosquitoes. ERL performed the CNV scan. HR provided intellectual input and drafted the manuscript.

