## Supplementary figures and images for "Multi-omics analysis identifies loci associated with pyrethroid resistance across sister species in the *Anopheles gambiae* species complex"

### Supplementary Figure 1

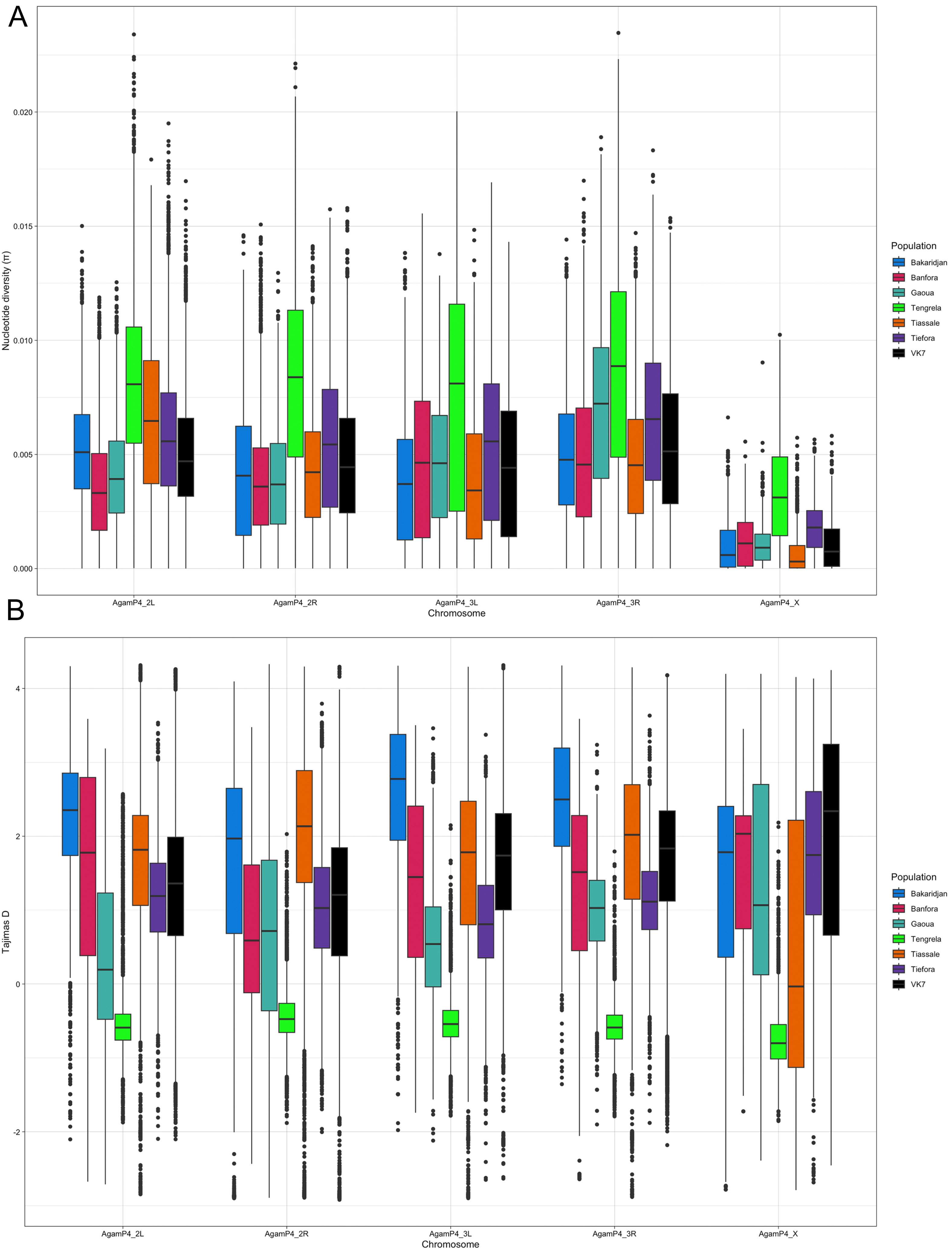

### Supplementary Figure 2

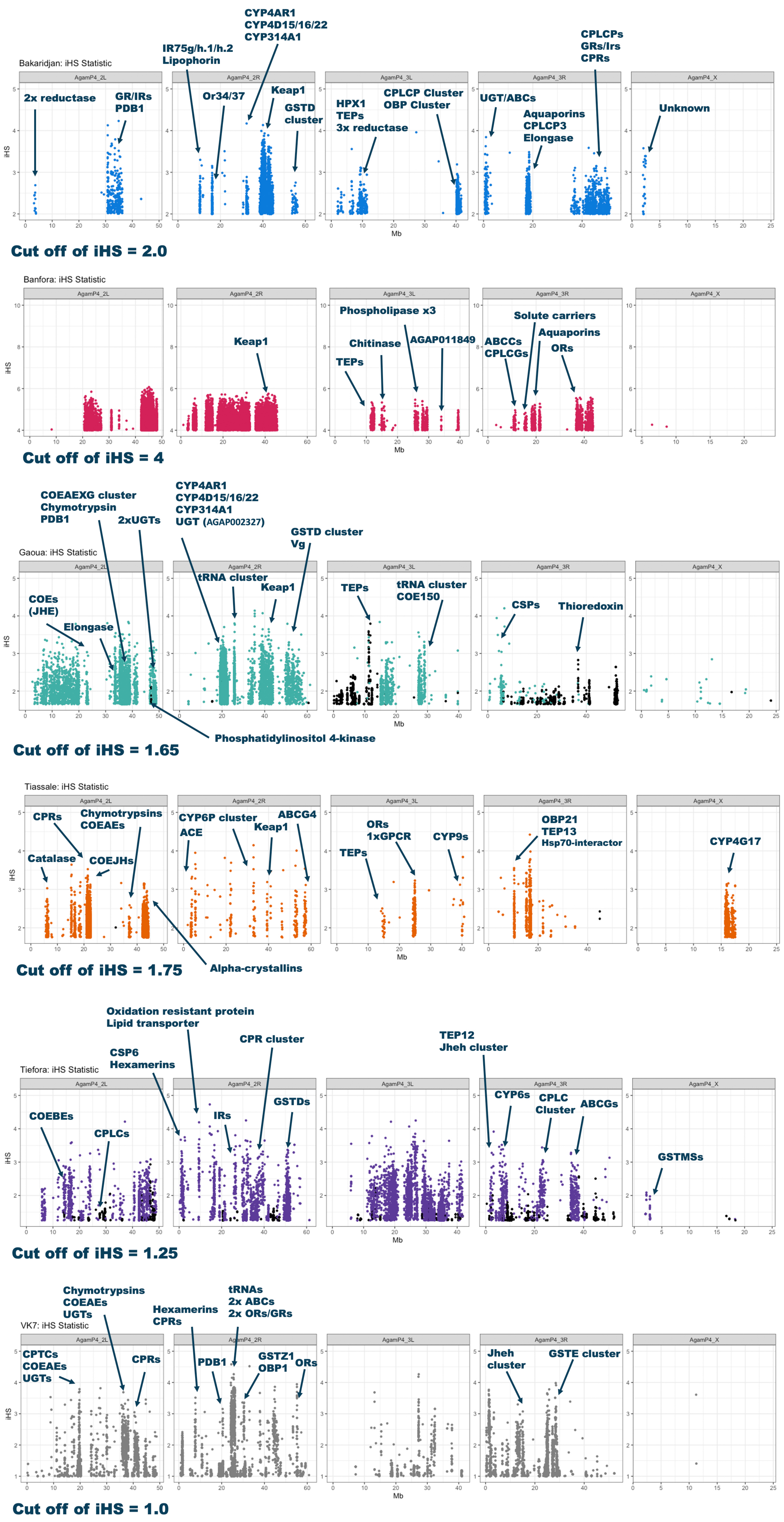

### Supplementary Figure 3

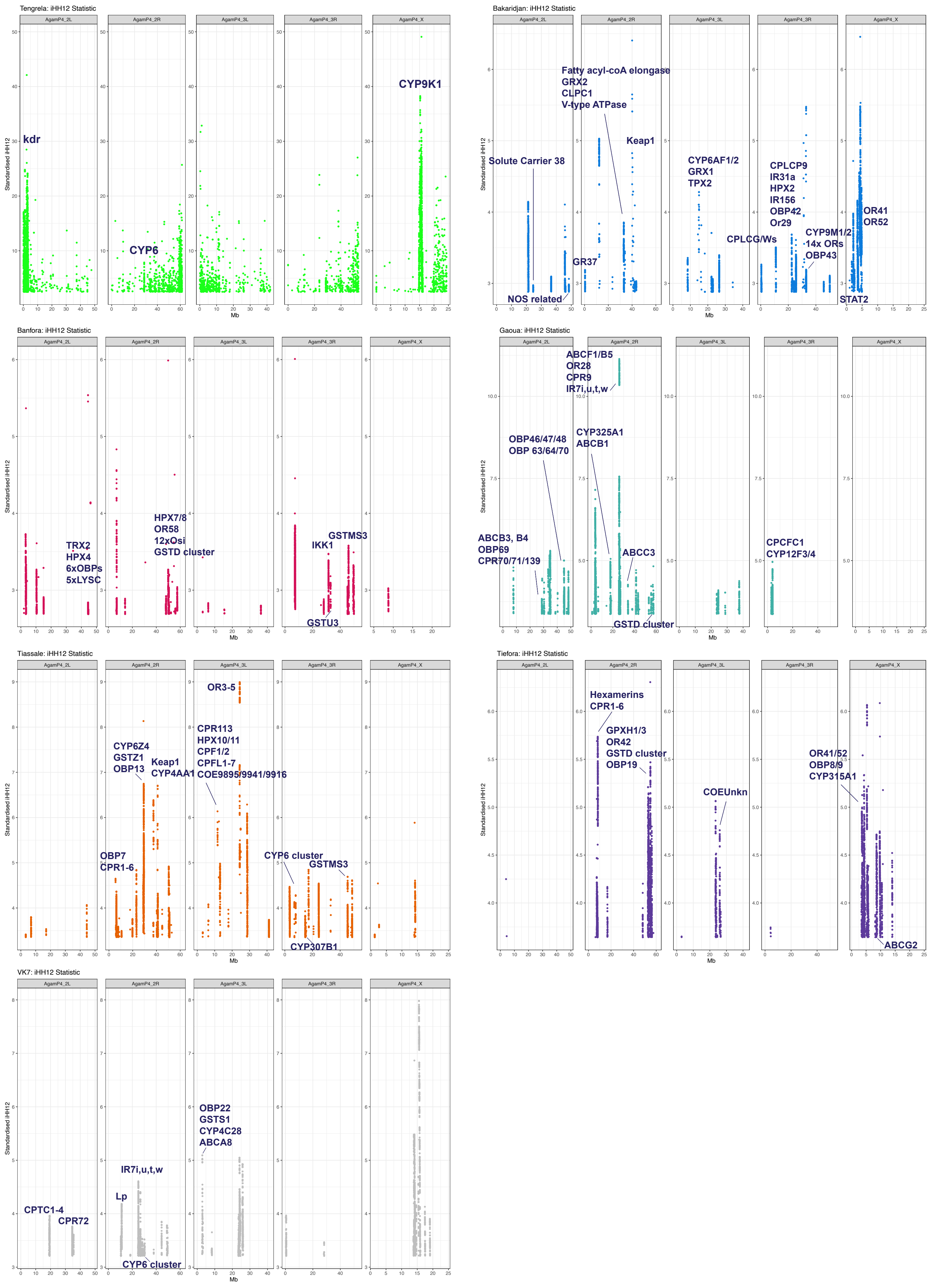

### Supplementary Figure 4

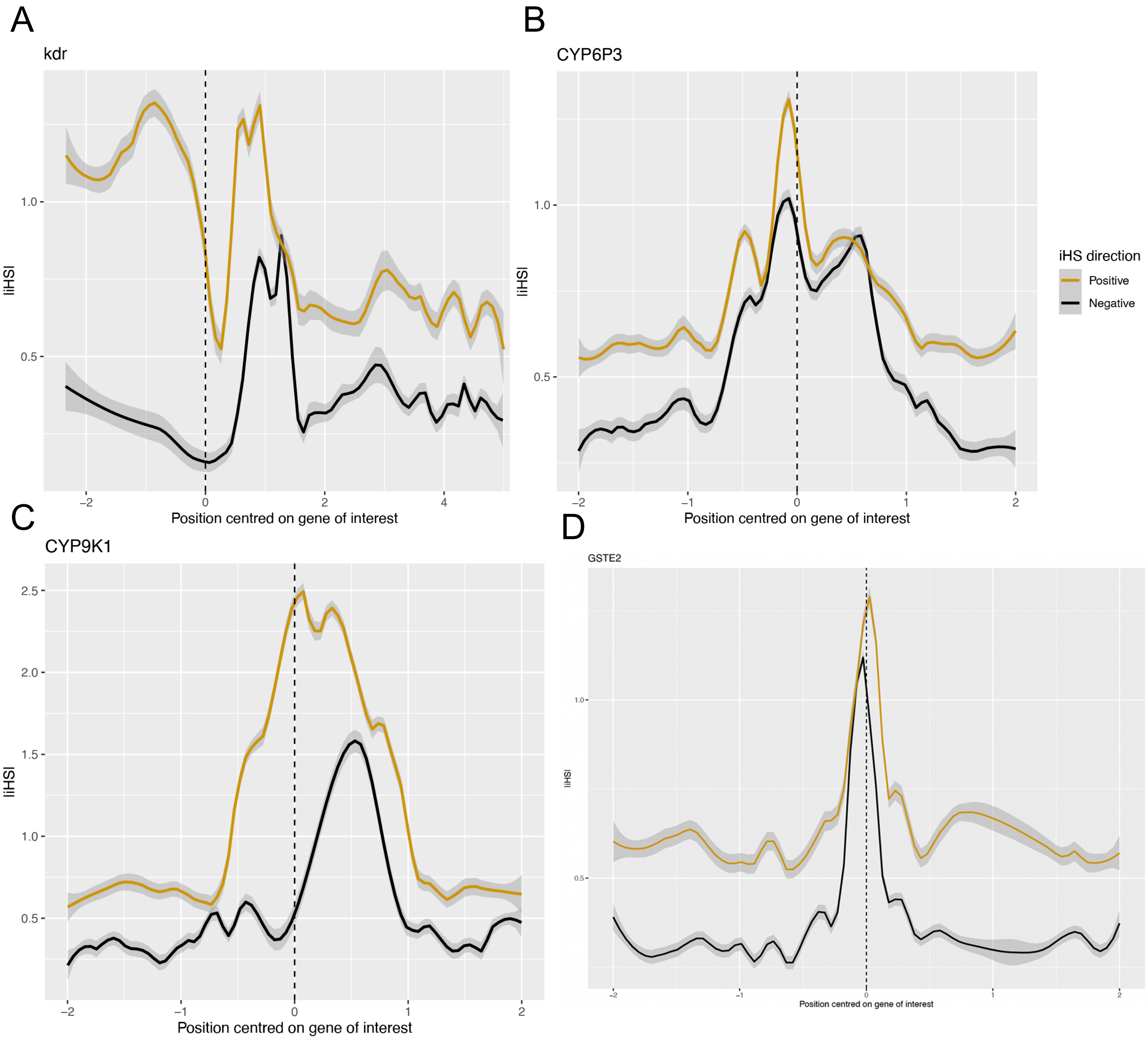

### Supplementary Figure 5

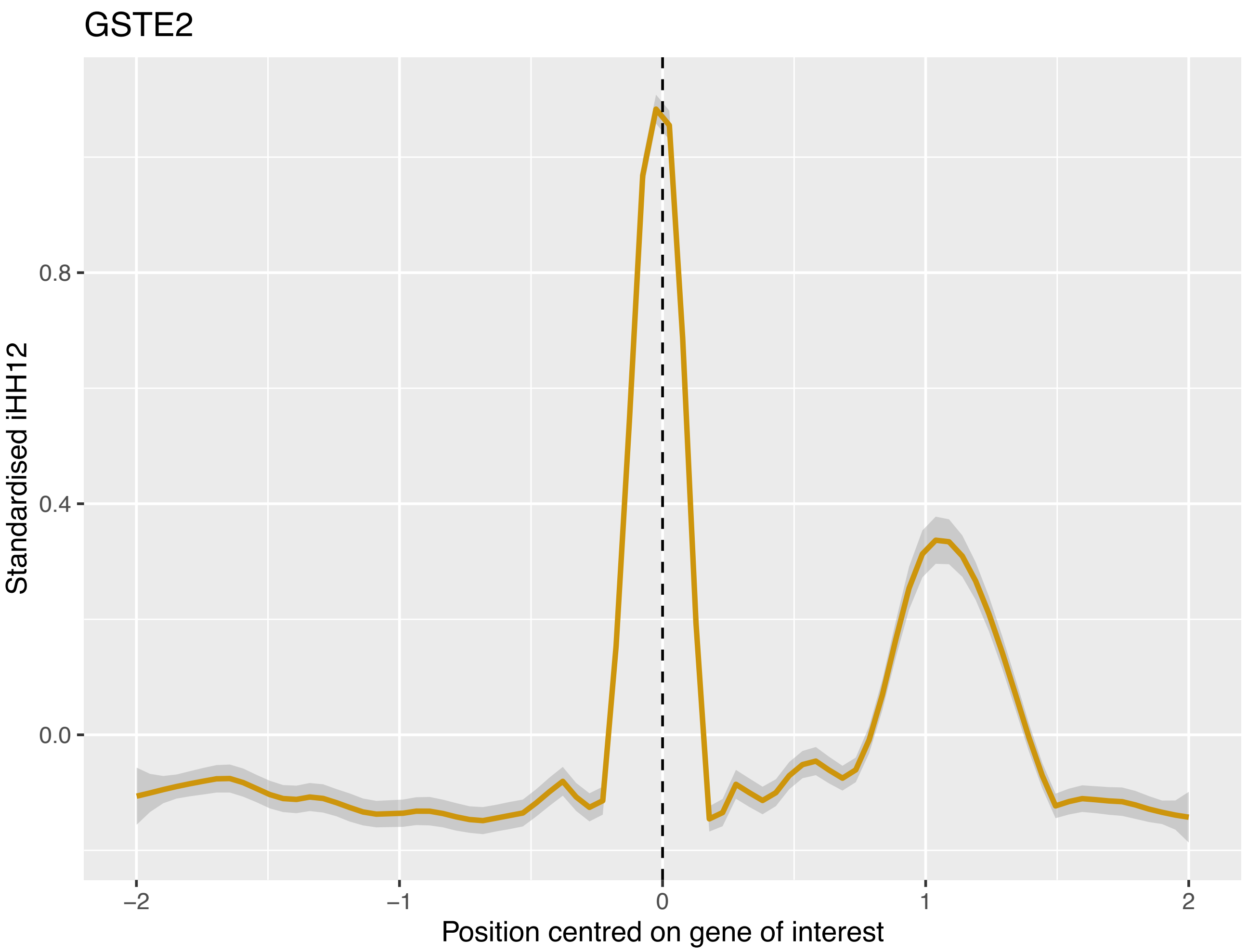

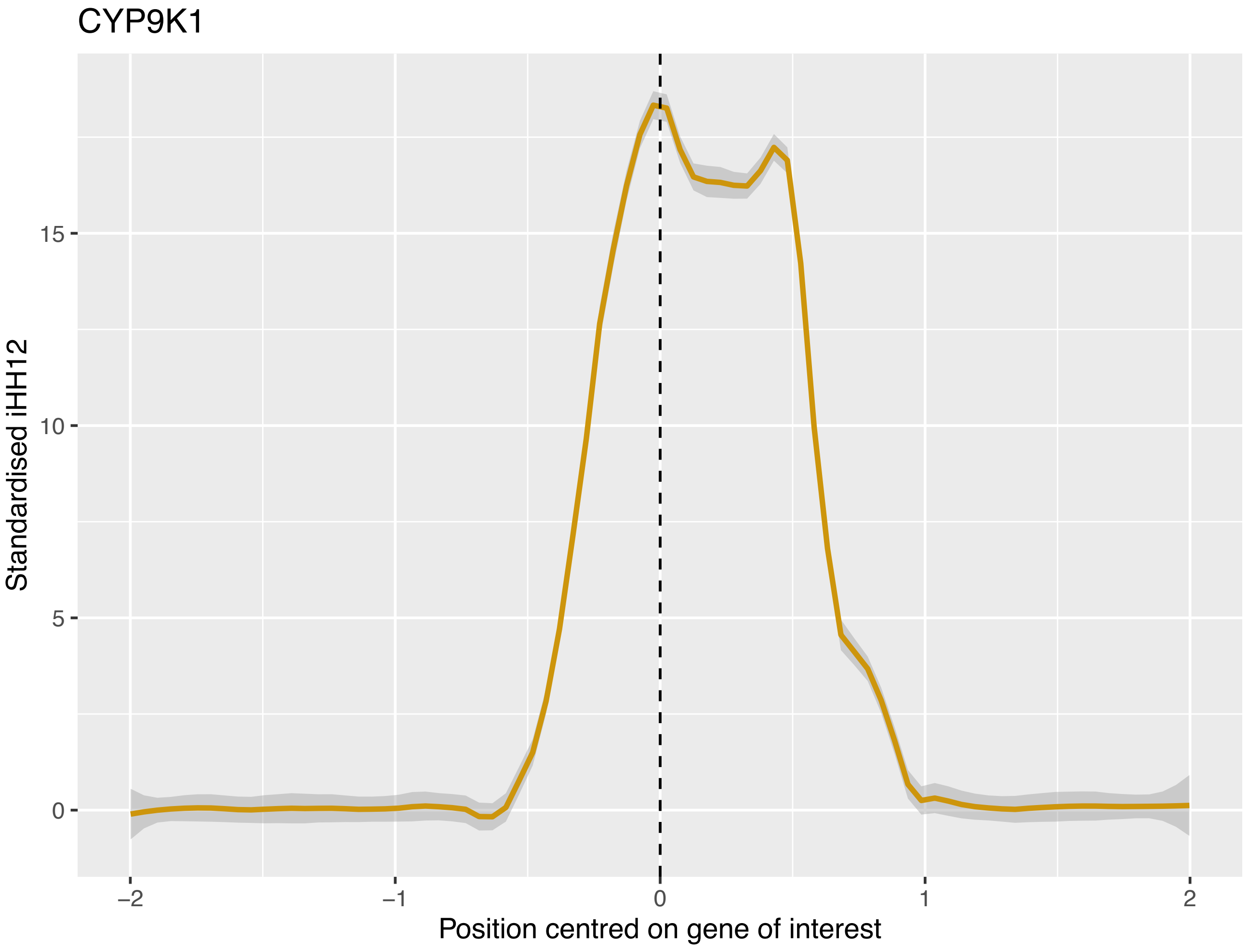

### Supplementary Figure 6

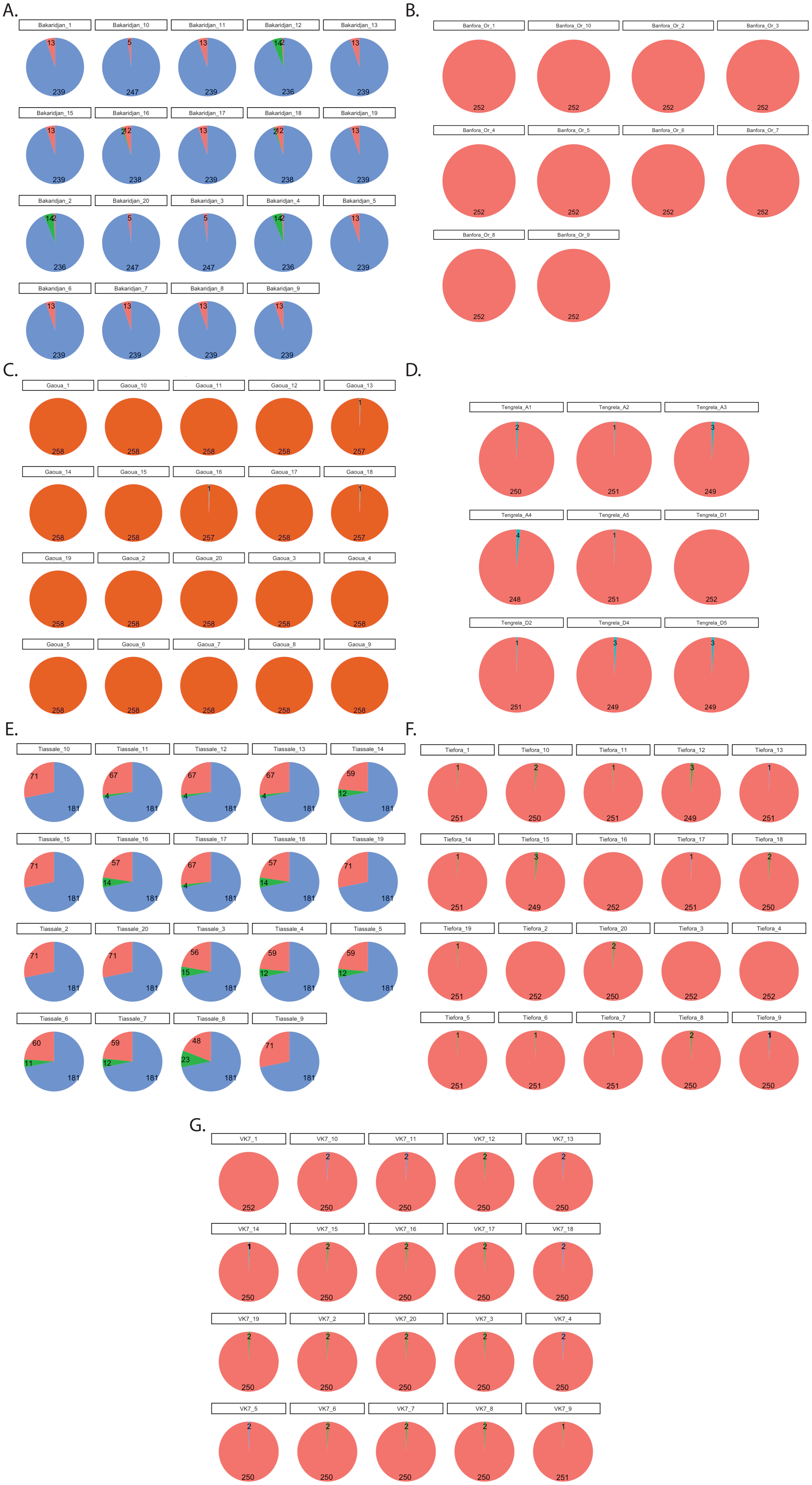

### Supplementary Figure 8

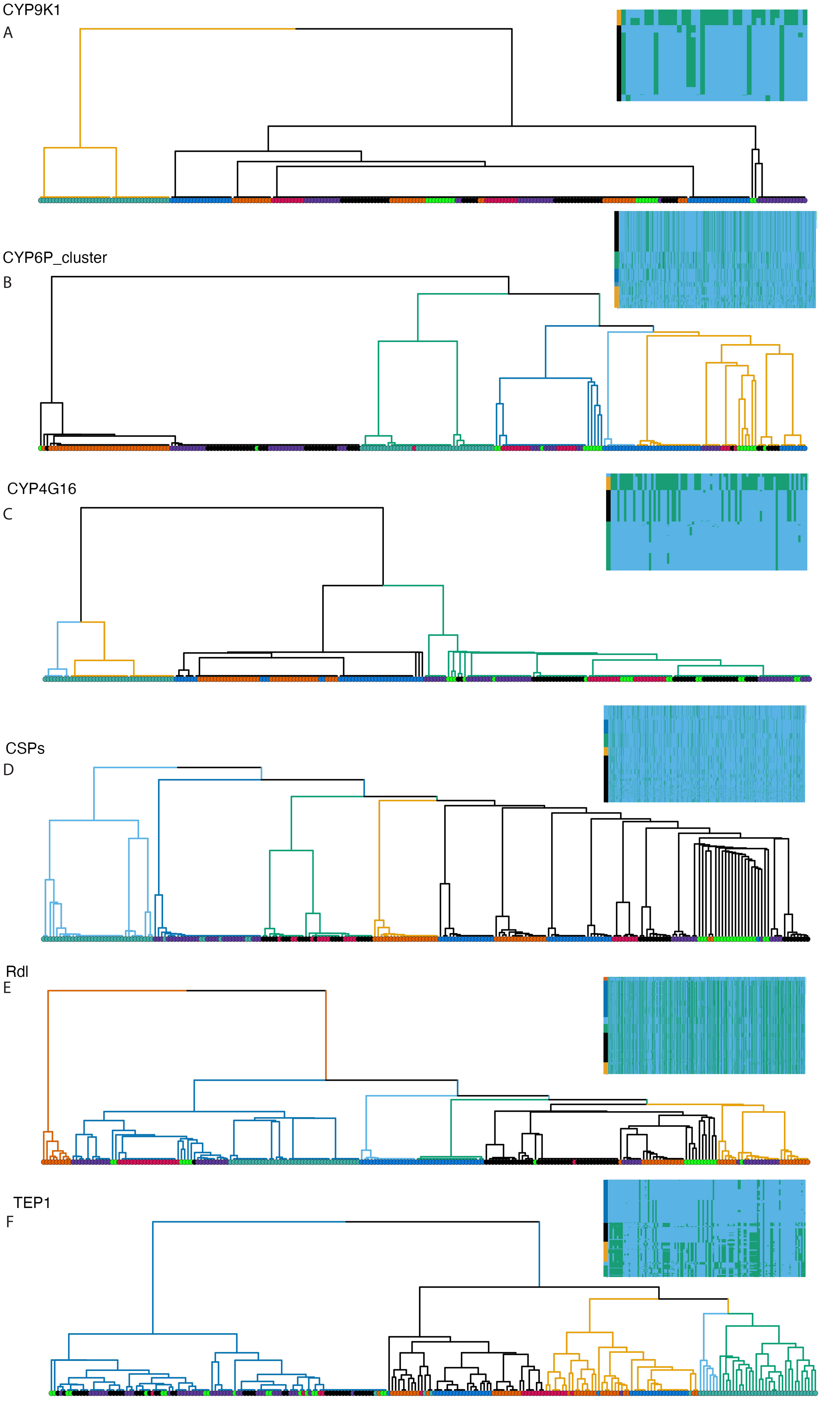

### Supplementary Figure 9

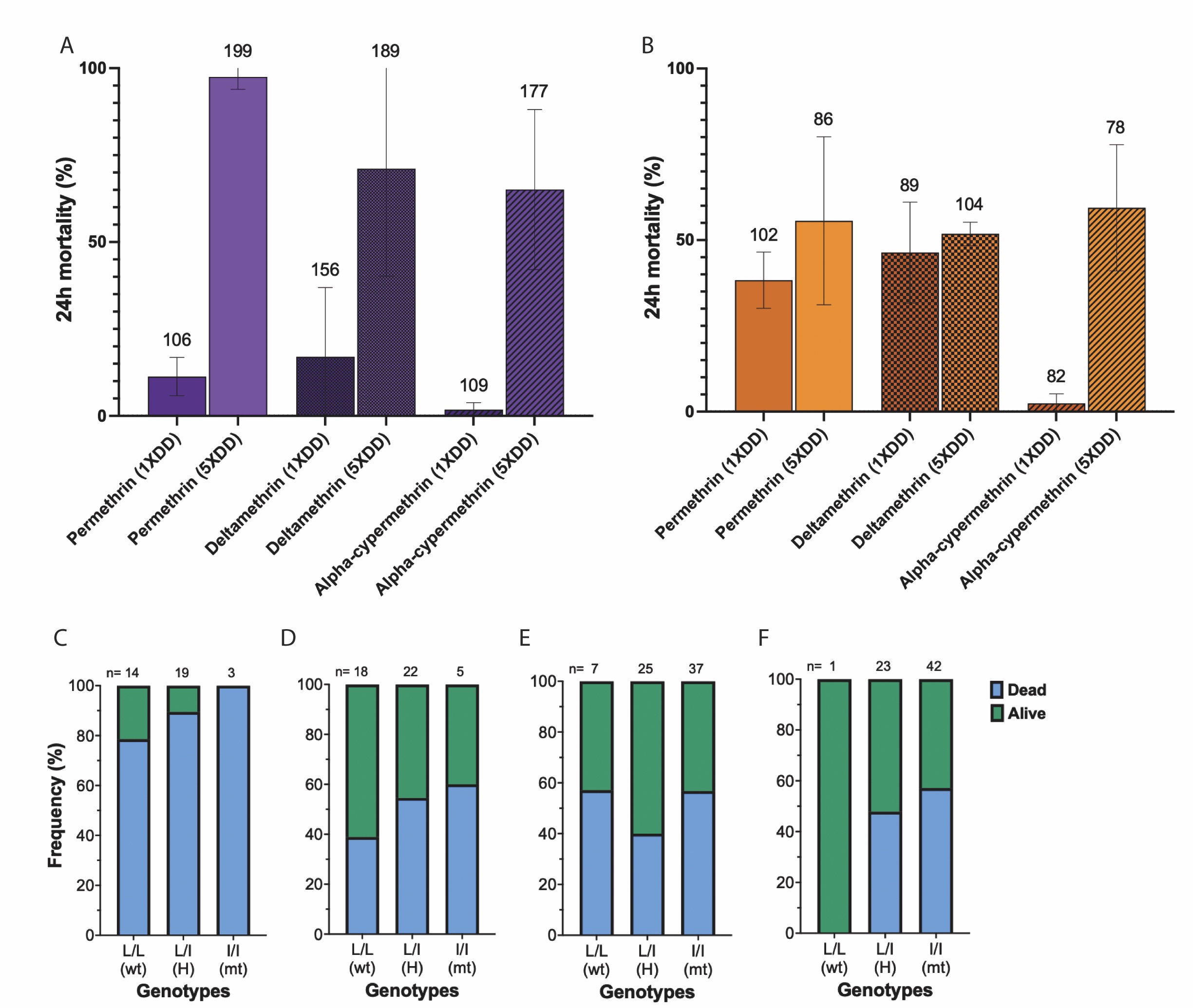

### Supplementary Figure 10

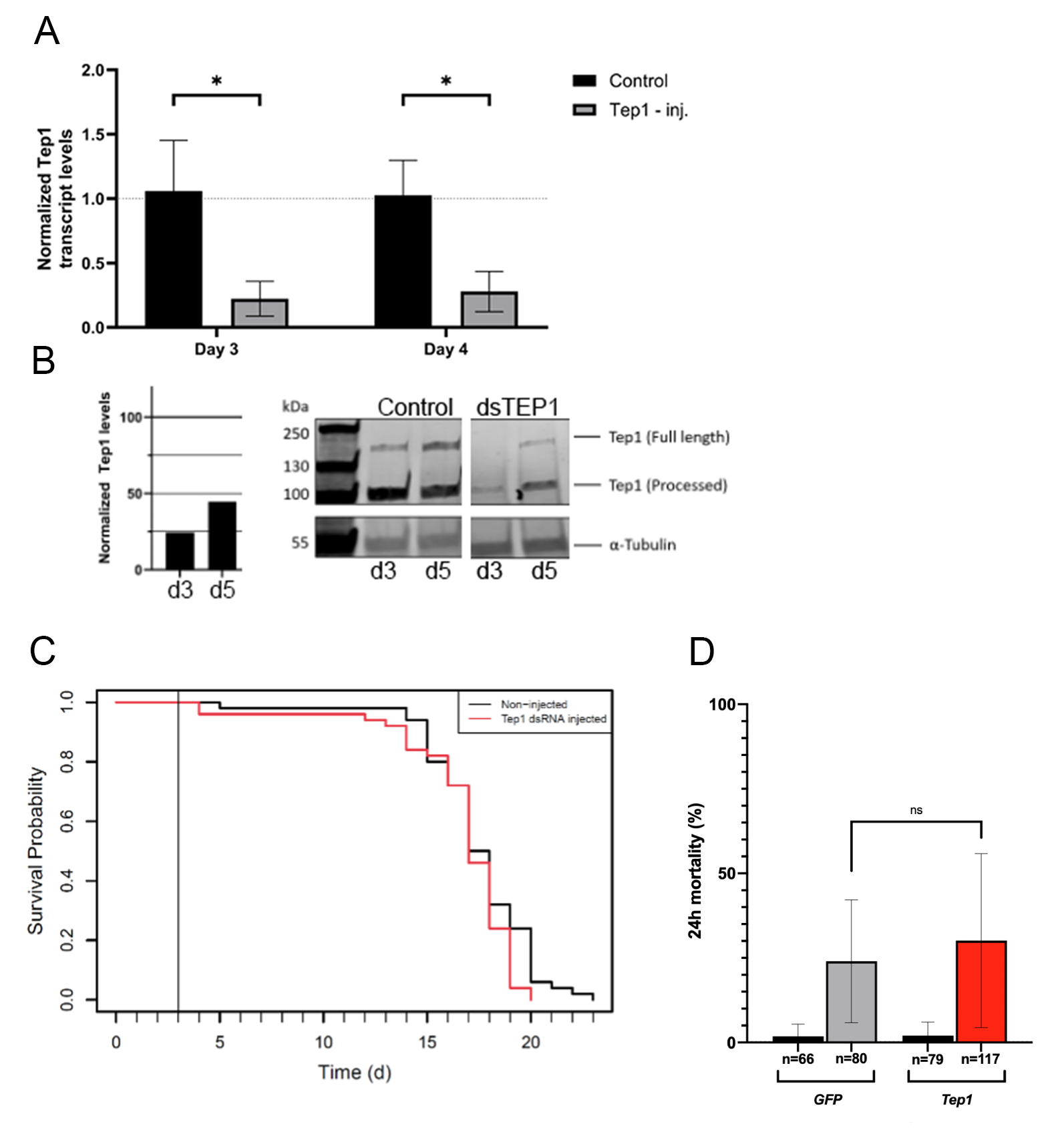

### Supplementary Figure 11

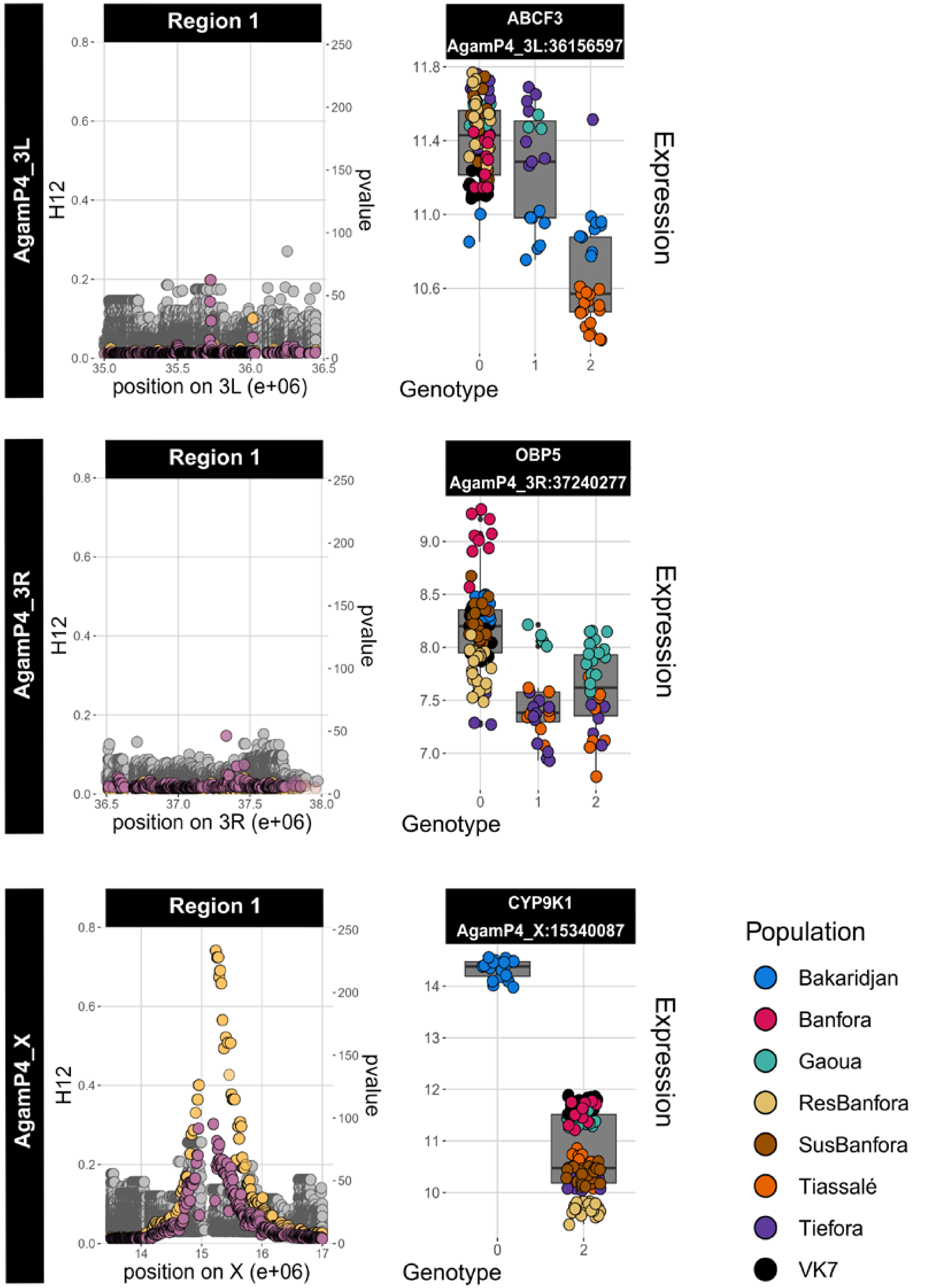

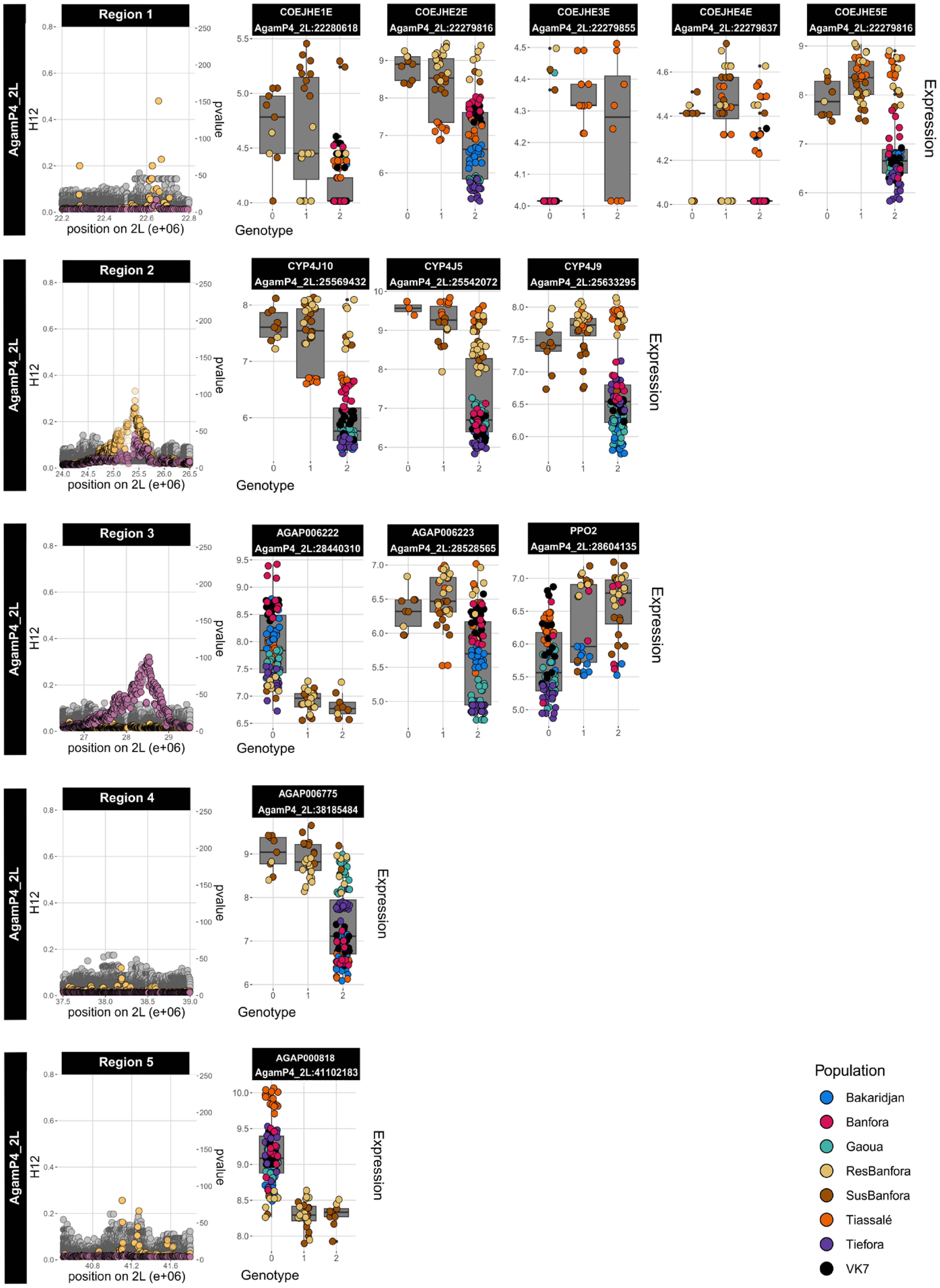

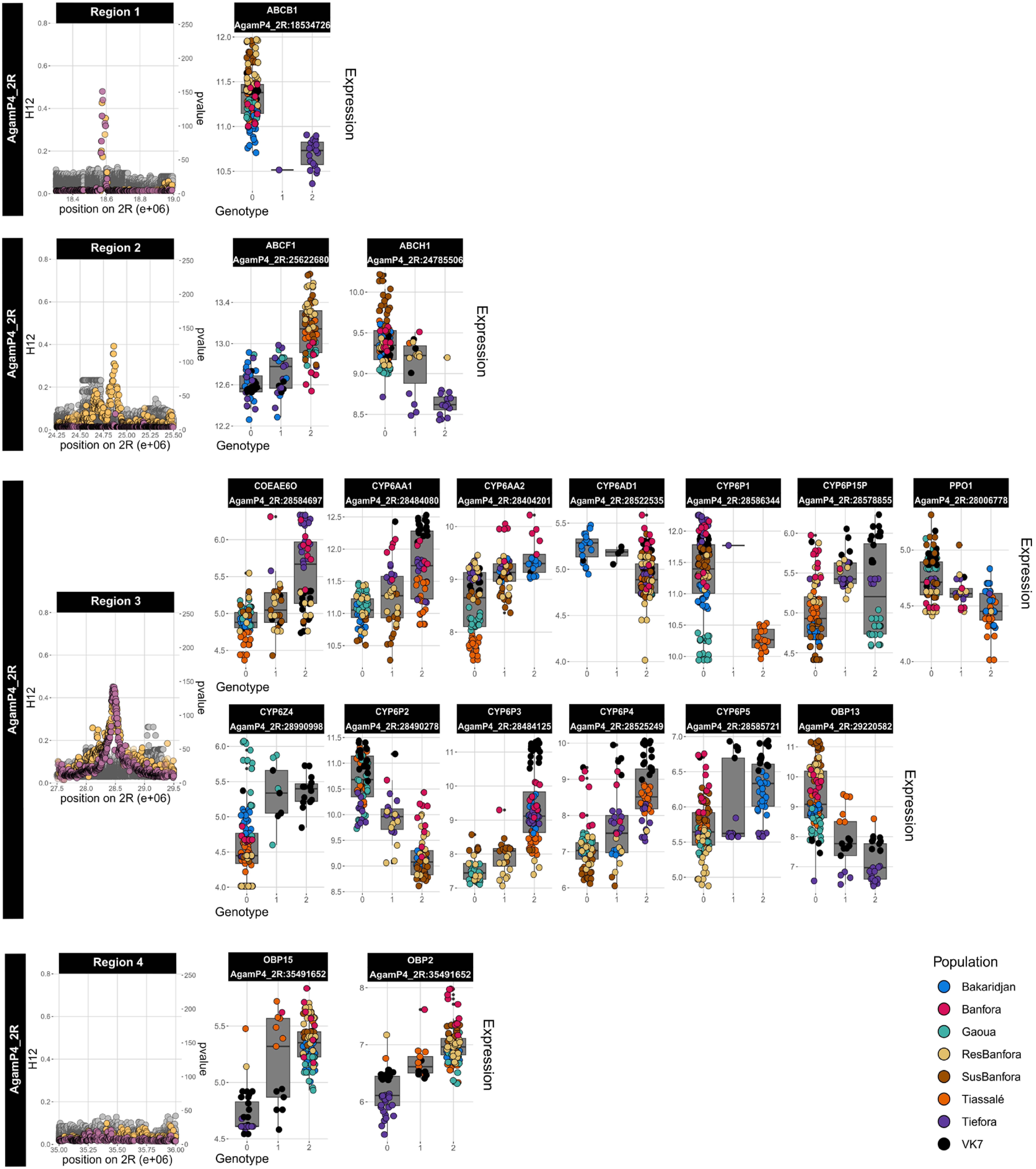
